# Modeling memory T cell states at single-cell resolution identifies *in vivo* state-dependence of eQTLs influencing disease

**DOI:** 10.1101/2021.07.29.454316

**Authors:** Aparna Nathan, Samira Asgari, Kazuyoshi Ishigaki, Tiffany Amariuta, Yang Luo, Jessica I. Beynor, Yuriy Baglaenko, Sara Suliman, Alkes Price, Leonid Lecca, Megan B. Murray, D. Branch Moody, Soumya Raychaudhuri

**Affiliations:** Center for Data Sciences, Brigham and Women’s Hospital and Harvard Medical School; Boston, MA 02115, USA; Division of Rheumatology, Inflammation, and Immunity, Department of Medicine, Brigham and Women’s Hospital and Harvard Medical School; Boston, MA 02115, USA; Division of Genetics, Department of Medicine, Brigham and Women’s Hospital and Harvard Medical School; Boston, MA 02115, USA; Program in Medical and Population Genetics, Broad Institute of MIT and Harvard; Cambridge, MA 02115, USA; Department of Biomedical Informatics, Harvard Medical School; Boston, MA 02115, USA; Department of Epidemiology, Harvard T. H. Chan School of Public Health; Boston, MA 02115, USA; Department of Biostatistics, Harvard T. H. Chan School of Public Health; Boston, MA 02115, USA; Department of Global Health and Social Medicine, Harvard Medical School; Boston, MA 02115, USA; Socios En Salud Sucursal; Peru, 15001, Lima, Peru; Division of Global Health Equity, Department of Medicine, Brigham and Women’s Hospital and Harvard Medical School; Boston, MA 02115, USA; Centre for Genetics and Genomics Versus Arthritis, Manchester Academic Health Science Centre, University of Manchester; Manchester M13 9PL, UK

## Abstract

Many non-coding genetic variants cause disease by modulating gene expression. However, identifying these expression quantitative trait loci (eQTLs) is complicated by gene-regulation differences between cell states. T cells, for example, have fluid, multifaceted functional states *in vivo* that cannot be modeled in eQTL studies that aggregate cells. Here, we modeled T cell states and eQTLs at single-cell resolution. Using >500,000 resting memory T cells from 259 Peruvians, we found over one-third of the 6,511 *cis*-eQTLs had state-dependent effects. By integrating single-cell RNA and surface protein measurements, we defined continuous cell states that explained more eQTL variation than discrete states like CD4+ or CD8+ T cells and could have opposing effects on independent eQTL variants in a locus. Autoimmune variants were enriched in cell-state-dependent eQTLs, such as a rheumatoid-arthritis variant near *ORMDL3* strongest in cytotoxic CD8+ T cells. These results argue that fine-grained cell state context is crucial to understanding disease-associated eQTLs.

---

Genome-wide association studies (GWAS) of autoimmune and allergic diseases have implicated non-coding variants that may regulate T cell gene expression (*1–5*). However, studies measuring the effect of these variants on bulk gene expression— expression quantitative trait loci (eQTL)—have incompletely explained their pathogenicity (*6*). Bulk assays obscure heterogeneity that is essential for effective T cell function and often require non-physiologic *ex vivo* stimulation. Single-cell assays, on the other hand, capture fine-grained physiologic T cell states defined by discrete surface markers (CD4+, CD8+), cytokines (T_H_1, T_H_2, T_H_17), transcription factors (T-bet, RORγt), or transcriptomic programs with varying degrees of expression (effector, cytotoxicity, activation). These states are neither static nor mutually exclusive: they may coexist in the same cell (for example, more effector-like CD4+ T_H_2 cells, seen in asthma) or they may transition (for example, T_H_17 cells gradually become IFNγ- and IL-17-coproducing T_H_17/1 cells seen in tuberculosis antigen-specific cells) (*7–10*). Certain states are effective therapeutic targets, like T_H_2s in allergy and T_H_17s in psoriasis (*11, 12*).

A T cell’s states may determine the magnitude or presence of eQTLs in that cell. For example, *ex vivo* activation alters variants’ regulatory effects (*13, 14*). However, most recent single-cell eQTL studies are unable to achieve this resolution because they identify state-dependent effects by first aggregating cells from each discrete cluster or other phenotypic classification to reduce dimensionality and mitigate sparsity and then using linear models (*15–18*). This limits the scope of analysis to coarse states that may imperfectly capture T cell biology or arbitrarily partition a continuous transcriptional landscape, such as single-cell differentiation trajectories or functional gradients like cytotoxicity.

In order to understand how regulatory genetic variants interact with the dynamic range of *in vivo* T cell states essential for disease pathogenesis, here we instead leverage multidimensional cell-state heterogeneity captured in multimodal single-cell assays of resting memory T cells. By considering each cell’s position along multiple continuous functional axes, we can dissect state-dependent eQTL effects at single-cell resolution and better identify disease-relevant regulatory heterogeneity.

## Results

### Memory T cell eQTLs in Peruvians include shared and ancestry-specific regulatory variation

For this study, we used single-cell expression of the transcriptome and 30 surface proteins from our previous CITE-seq study of memory T cells isolated from 259 healthy Peruvian individuals with prior resolved *Mycobacterium tuberculosis* infection (*10*). We chose the surface proteins because of their role in T cell function. After applying quality control (QC) as shown previously, 500,089 cells remained for eQTL analysis, with an average of 4,927 unique molecular identifiers (UMIs) and 1,475 genes per cell (**Methods**, **Fig. S1A-B**) (*10*). We analyzed 5,460,354 genotyped and imputed variants passing QC.

We first defined a core set of eQTLs across all memory T cells, agnostic to state; eQTLs demonstrating a robust main effect would be promising candidates to later test for state-dependent effects in a single-cell model (**Fig. 1A**). Since we were not yet considering individual cells’ states, pseudobulk analysis was sufficient. We summed the expression of each gene across all cells from each donor (mean = 1,851 cells/donor, **Fig. S1C**) and treated this expression profile as a bulk sample for sample-level normalization, gene QC, and correction of measured and latent covariates. Next, to define *cis*-eQTL effects, we tested associations between the covariate-corrected expression of 15,789 genes expressed in >50% of samples and all variants up to 1 MB from each gene’s transcription start site (TSS).

**Fig. 1.**
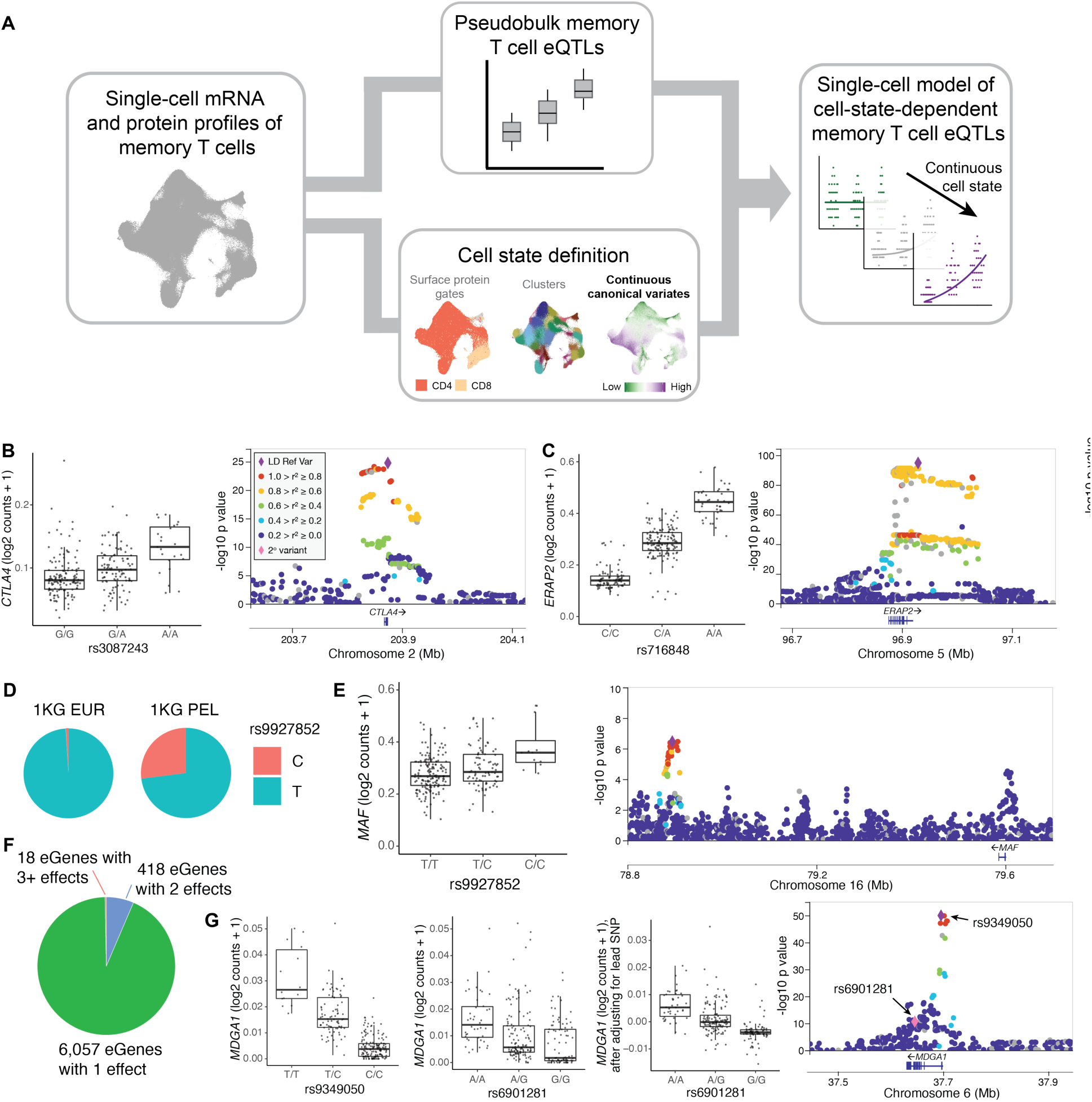
Modeling memory T cell eQTLs. **(A)** Schematic of single-cell eQTL modeling strategy. Using single-cell CITE-seq profiles, we conduct pseudobulk eQTL analysis to define memory T cell eQTLs and single-cell states. Specifically, continuous canonical variates can be used to dissect state-dependence of the memory T cell eQTLs in a single-cell model (shown here binned into low/medium/high for ease of visualization). **(B)** (left) Box plot and (right) locus plot of rs3087243 eQTL for *CTLA4*. Except where indicated, each point in a box plot represents the average log2(UMI counts + 1) across all cells in a donor (n = 259), grouped by genotype. Box plots show median (horizontal bar), 25th and 75th percentiles (lower and upper bounds of the box, respectively) and 1.5 times the IQR (or minimum/maximum values if they fall within that range; end of whiskers). Each locus plot shows the variants in a +/250kb window around the TSS plotted based on their nominal pseudobulk eQTL p value and genomic coordinate. The purple diamond is the lead variant and other variants are colored based on their r^2^ with the lead variant. **(C)** Box plot and locus plot of rs716848 eQTL for *ERAP2*. **(D)** Pie charts of the allele frequencies at rs99278652 in 1KG EUR (European) and PEL (Peruvian in Lima) populations. **(E)** Box plot and locus plot of rs9927852 eQTL for *MAF*. **(F)** Number of eGenes with 1, 2, or 3+ independent eQTLs. **(G)** Box plots for lead (rs9349050, left) secondary (rs6901281, middle), and secondary conditioned on lead (right) eQTL variants for *MDGA1*. In the box plot for rs6901281 conditioned on rs9349050, each point represents the average residual of log2(UMI counts + 1) after regressing out genotype at rs9349050 across all cells in a donor (n = 259). In the locus plot, the pink diamond is the secondary variant.

We found 6,511 eGenes with significant *cis*-eQTLs (q value < 0.05), consistent with previous bulk eQTL studies with similar sample size (*19, 20*). These genes included previously described eGenes such as *CTLA4* and *ERAP2* (**Fig. 1B-C, Table S1**) (*20, 21*). We also found 808 eQTLs that were driven by genetic variation common in the Peruvian population but rare or absent in Europeans **(Table S2)**. For example, an eQTL for *MAF* (β = 0.32, p = 3.45×10^-7^) was driven by rs9927852 (chr16_78894778_T_C: minor allele frequency=22% in study cohort, 27% in 1000 Genomes Peruvians in Lima, Peru [PEL], 1% in European [EUR], **Fig. 1D-E**) (*22*). When we conditioned on the lead eQTL for each eGene (n = 6,511), we observed exactly two independent effects at 418 loci, such as *MDGA1*, and more than two independent effects at 18 loci upon repeated conditional analysis (**Fig. 1F-G, Table S3**).

To determine if results were consistent with previously published T cell eQTLs, we compared the lead variants’ effects to bulk naive CD4+ T cell eQTLs from individuals of European ancestry (n = 169) reported by the BLUEPRINT project (*19*). eQTL dynamics described in prior studies suggest that naive and memory T cells share many bulk eQTL effects (*2, 21*). Despite differences in linkage disequilibrium due to ancestry, technology, and cell type, we observed that the eQTLs from our analysis were largely significant in BLUEPRINT (at q < 0.05, 2,056 significant in both/3,249 significant in current study and measured in BLUEPRINT), and of those that were significant in both datasets, most had concordant directions of effect (1,917/2,056 = 93% same direction, **Fig. S2**).

### Continuous cell states capture functionally distinct dimensions of T cell heterogeneity

Combined single-cell mRNA and protein measurements from CITE-seq allow us to define cell states in conventional ways, such as clustering or gating on protein markers as in flow cytometry (e.g. CD4+ T cells). However, to better capture the continuous heterogeneity of T cell states, we projected cells into a multimodal low-dimensional embedding with canonical correlation analysis (CCA), as previously described (**Methods**) (*10*). Each cell received a score along 20 dimensions (canonical variates, CV) defined by orthogonal variation shared between mRNA and surface protein expression, because cross-modality signal is likely to reflect more robust cell states (**Fig. S3A**). This was demonstrated when we clustered on these CVs in the original study and identified 31 canonical memory T cell states including regulatory, type 1/2/17 helper, and gamma delta T cells, many of which could not be precisely defined in unimodal analysis of mRNA alone **(Fig. S3B)** (*10*).

Rather than using CVs to partition cells into clusters, as in the original study, we now used the top CVs as continuous representations of cell state (**Fig. 1A**). We expected that each CV might represent a distinct, biologically relevant function because clusters delineated by the CVs correspond to known T cell states (**Fig. S3B**). Indeed, we observed that individual CVs correlate with genes, proteins, and gene sets relevant to well-described T cell functions (**Fig. 2A**). For example, CV1 correlated with a previously defined cytotoxicity (“innateness”) gene set and CV2 correlated with a regulatory T cell (T_reg_) gene set (**Fig. 2B-C**) (*23, 24*). We confirmed both correlations with gene set enrichment analysis (**Table S4**). In some instances, published gene sets weren’t significantly enriched, but CVs correlated with marker genes of known memory T cell states. For example, CV4 correlated with T_H_2 marker *GATA3* (Pearson r = 0.23 in non-zero cells, p < 10^-1785^), and CV8 correlated with gamma delta T cell marker *TRDC* (Pearson r = 0.51 in non-zero cells, p < 10^-767^; **Fig. 2A, Fig. S3C-F**) (*25*).

**Fig. 2.**
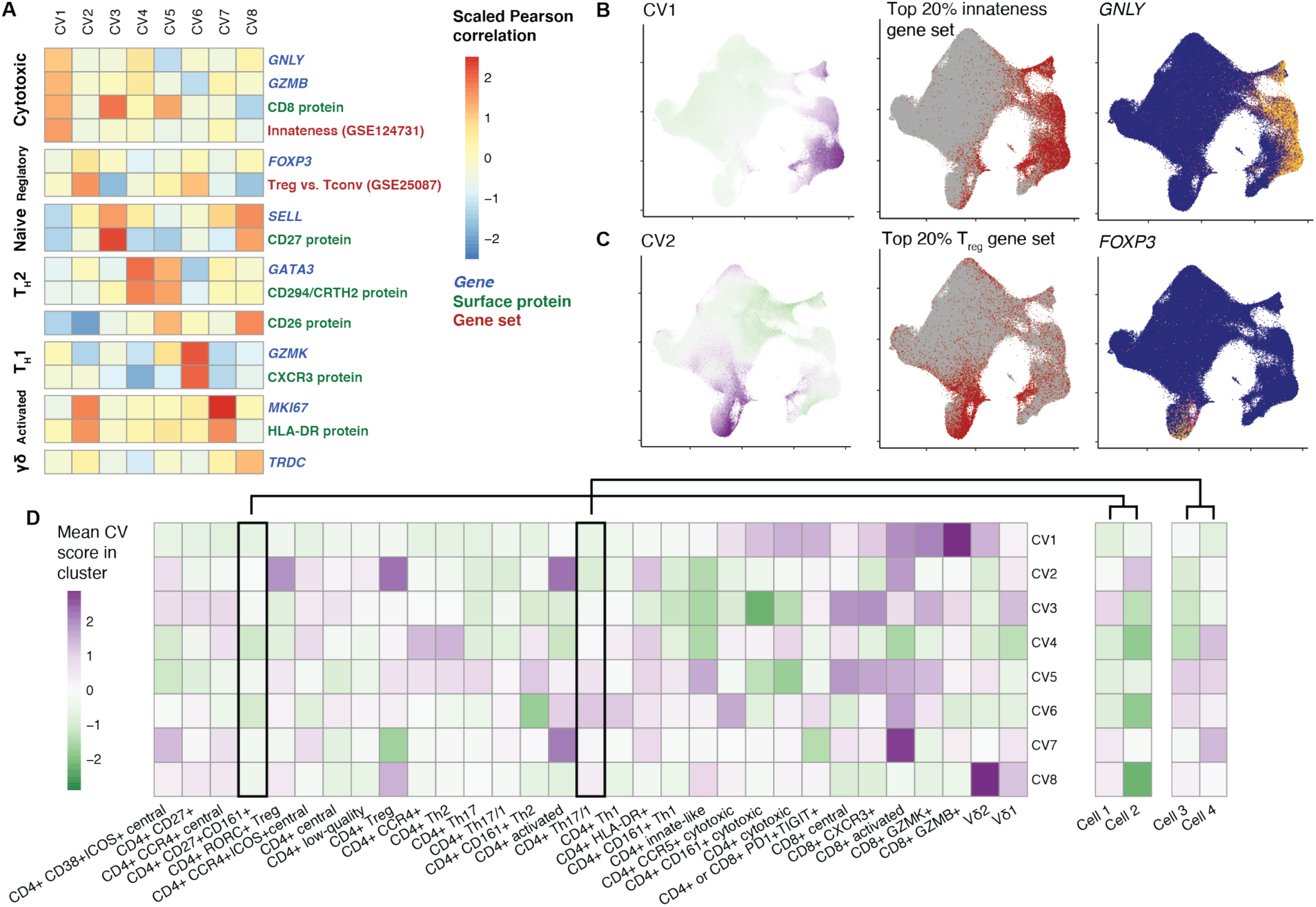
Canonical variates (CVs) reflect biological functions. **(A)** Heatmap colored by scaled Pearson correlations between CVs and select marker genes, surface proteins, and gene set scores. Correlations were computed with log2(counts per 10,000)-normalized expression for genes, centered-log-ratio normalized expression for proteins, and summed normalized expression of genes in the gene set (weighted where available). **(B)** UMAPs of memory T cells colored by CV1 score (left), top 20% of cells based on summed weighted expression of innateness gene set (in red), and normalized expression of *GNLY*. **(C)** UMAPs of memory T cells colored by CV2 score (left), top 20% of cells based on summed expression of T_reg_ gene set (in red), and normalized expression of *FOXP3*. Colors for CV scores range from low (green) to high (purple) for each CV. Colors for gene expression range from minimum (blue) to maximum (yellow) for each gene. **(D)** Heatmap colored by average score for each CV (1-8) among cells in each cluster. Separate columns on the right show CV scores for two cells each from two clusters. Colors range from low (green) to high (purple).

Single-cell analyses typically use multiple components of a low-dimensional embedding to define higher-resolution cell states—often clusters—carrying out combinations of functional programs (*26, 27*). Accordingly, the average CV scores of T cells in each CCA-defined cluster varied (**Fig. 2C, Table S5**) and can be used to interpret the functional diversity among clusters, e.g., some clusters have more effector function (high CV1), others are more T_H_1-like (high CV6). However, clusters obscure heterogeneity that exists between individual cells from the same cluster **(Fig. 2C)**. Moreover, continuous metrics like CVs don’t just classify cells into states, but instead capture the degree of how much each state influences a cell, which is a more faithful representation of how activation or helper states manifest in T cells (*8, 28*). Thus, using the CV scores themselves—or similar metrics defined at single-cell-resolution—may be more precise.

### Single-cell statistical models define cell-state-dependent eQTLs

Modeling sparse expression and cell states at single-cell resolution requires statistical models that differ from those commonly used for bulk or pseudobulk eQTL analysis. Here, we used single-cell Poisson mixed-effects (PME) regression, which can model discrete and continuous cell states, Poisson-distributed scRNA-seq UMI counts, and the nested structure of cells within donors and batches (*29, 30*). We model the UMI counts of each gene in single cells as a function of genotype, correcting for potentially confounding fixed-effect covariates (age, sex, genotype PCs, gene expression PCs) and random-effect covariates (donor, batch) (**Fig. 3A, Methods**).

**Fig. 3.**
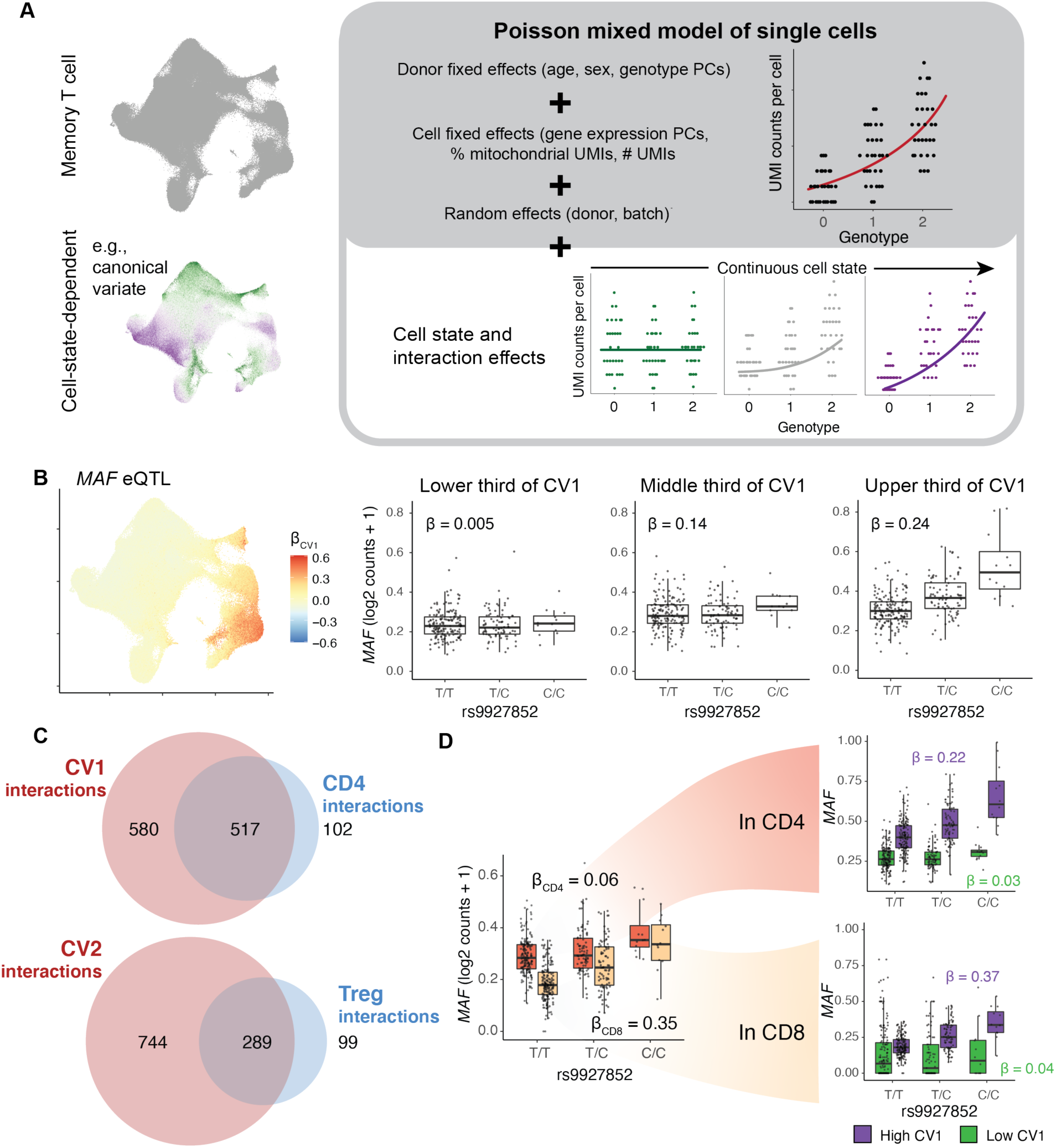
Modeling eQTL interactions with continuous cell states across single cells. **(A)** Schematic of Poisson mixed effects model for cell-state-dependent single-cell eQTL analysis. After adjusting for covariates and modeling random effects, we can measure the interaction between a continuous cell state (shown here binned into low/medium/high for ease of visualization) and genotype. **(B)** Interaction of rs9927852 eQTL for *MAF* with CV1. UMAP of total effect size (β_total_ = β_G_ + β_CV1_*CV1 score) per cell. Box plots show eQTL effect for cells in the bottom (left), middle (center), and top (right) thirds of CV1 scores. **(C)** Venn diagrams of the number of eGenes with significant CV interactions (red) and discrete state/cluster interactions (blue) for CV1 compared to CD4+ (top) and CV2 compared to T_reg_ (bottom). eGenes interacting with both the CV and the discrete state are indicated in the overlap. Interactions are deemed significant at q < 0.05. **(D)** rs9927852 eQTL for *MAF* in subsets of cells. The box plot on the left shows the eQTL in CD4+ (orange) and CD8+ (beige) cells. The box plots on the right show the eQTL in CD4+ cells (top) and CD8+ cells (bottom) divided by low (green) or high (purple) CV1 score. CD4+ or CD8+ classification is based on CITE-seq surface-protein-based gating of TCRab+CD4+CD8- and TCRab+CD4-CD8+ cells. CV1 high or low is based on threshold = 0. For **(B)** and **(D),** each point in a box plot represents the average log2(UMI counts + 1) across all cells in the indicated subset of cells in a donor (n = 259), grouped by genotype. Box plots show median (horizontal bar), 25th and 75th percentiles (lower and upper bounds of the box, respectively) and 1.5 times the IQR (or minimum/maximum values if they fall within that range; end of whiskers).

To demonstrate consistency with commonly used linear models, we used the PME model to reanalyze our data and successfully recapitulated almost all eQTLs detected in pseudobulk analysis with nominal significance (6,291/6,511=97%) and concordant direction of effect (6,509/6,511=100%, **Fig. S4A, Table S6**). We permuted genotypes and observed that in this null data, 5.3% (347/6,511) of the eGenes were significant at p<0.05, demonstrating well-calibrated type 1 error. (**Fig. S4B**). Although donors were part of a former TB progression cohort, 6,510/6,511 eQTLs had no significant differences (q<0.05) between people with and without a history of disease progression (**Table S7**).

Then, to identify eQTLs with cell-state-dependent effects, we added an interaction term between genotype and cell state (capturing heterogeneity in the eQTL effect) to the PME model. We compared this model to a baseline model controlling for the genotype (overall eQTL effect) and cell state (differential expression) to assess significance (**Methods, Fig. 2A**). Although this model can accommodate continuous states, in order to compare it to conventional pseudobulk models, we first chose a simple binary test case: CD4+ vs. CD4-, based on surface protein expression measured with CITE-seq (**Fig. S5A-D**). The total eQTL effect estimated in CD4+ cells (β_total_ = β_G_ + β_GxCD4_) with the PME interaction model was consistent with eQTL analysis with a pseudobulk linear model or a single-cell PME model of CD4+ T cells gated from the CITE-seq dataset (**Fig. S5E-F, Table S8-10**). Furthermore, using genotype permutations, we demonstrated that type I error for the interaction term was well-controlled at α=0.05 (397/6511 = 0.061, **Fig. S5G**).

An alternative is to apply a linear mixed-effects (LME) model to normalized single-cell expression data. Without considering cell state, LME performed similarly to PME (**Fig. S4C-E, Table S11**). However, when we added an interaction term, the LME model was not robust to differential expression between cell states. Even when the only difference between an eQTL’s effect in CD4+ and CD4- cells was due to artificially simulated differential expression, the LME model spuriously detected highly significant state-specific eQTLs, while the PME model did not (**Methods, Fig. S6**). This is consistent with previous studies showing that LME models inadequately describe the distribution of single-cell expression data (*29, 30*).

### eQTL effects vary systematically along continuous single-cell states

With the single-cell resolution of the PME model, we were able to demonstrate how eQTLs varied across continuous T cell states. We represented cell states with cells’ projections on the individual CVs defined in the original study (**Fig. 3A, Fig. S3A**) (*10*). We found that a large proportion of eQTLs are modified by these cell states. Focusing on CV1, which captured cytotoxic function, we observed that 1,097 of 6,511 memory T cell eQTLs had a significant interaction (q < 0.05, **Table S12**), i.e. the magnitude of the eQTL effect varies in cells depending on their CV1 score. For example, the rs9927852 eQTL for *MAF* had an interaction effect that amplified the eQTL in cells with higher CV1 scores (β_G_ = 0.098, β_GxCV1_ = 0.13). This means the eQTL has almost no effect in cells in the lower third of CV1 scores, but increased to maximum effect in the upper third of CV1 scores (Average β_total_ in lower third =0.005, average β_total_ in upper third = 0.24, **Fig. 3B**). Interaction effects were independent from differential expression and main genotype effects, and the type 1 error was well-controlled upon permutation of CV1 scores (**Fig. S7**).

We observed that continuous cell states captured more state-dependent regulatory variation than analogous discrete phenotypes. For example, CD4+ and CD8+ are two major discrete lineages of memory T cells, and continuous CV1 scores largely discriminate between them (classifying CD4+ based on CV1 < 0: sensitivity = 0.85, specificity = 0.93; **Fig. S8A**). A PME model of eQTL interactions with CV1 recapitulated 517/619 (84%) eQTLs identified in a PME model with the CD4+ state, as expected, but also identified an additional 580 eQTLs uniquely significant in the continuous analysis (**Fig. 3C, Fig. S8B**). These eQTL interactions’ directions of effect were consistent between CV1 and CD4+, but they were non-significant in the discrete CD4+ analysis (94% concordant effect direction). Similarly, CV2 correlates with T_reg_ markers, and the 1,033 eQTLs with CV2 interactions included but exceeded the 289/388 (74%) eQTLs with significant T_reg_ cluster interaction effects (**Fig. 3C, Fig. S8C, Table S13**). These correspondences were specific, i.e., the CV1 eQTL interactions were not concordant with T_reg_ cluster interactions and CV2 eQTL interactions were not concordant with CD4+ gate interactions (**Fig. S8D**). This shows that the continuous states captured by decomposing single-cell data represent biological programs with regulatory significance.

Continuous cell states may also explain heterogeneous eQTL effects better than discrete states do. For example, the *MAF* eQTL interacts with both CD4+ status (β_G_ = 0.33, β_GxCD4_ = -0.25, p_GxCD4_ = 4.69×10^-85^) and CV1 (β_G_ = 0.098, β_GxCD4_ = 0.13, p_GxCD4_ = 1.80×10^-243^) in a biologically concordant manner, i.e., CD4+ cells tend to have lower CV1 scores and both are associated with weaker *MAF* eQTL. However, the interaction with CD4+ status was no longer significant when we conditioned on CV1 interactions (p = 0.77), but the CV1 interaction maintained significance (p = 1.19 x 10^-231^). Closer inspection of minor subsets of CD4+ T cells with higher CV1 scores (15%) and CD8+ T cells with lower CV1 scores (4.7%) confirms this. Both CD4+ and CD8+ memory T cells with high CV1 scores have strong eQTL effects (β_CD4,high CV1_ = 0.22, β_CD8,high CV1_ = 0.37, **Fig. 3D**), while cells of both lineages with low CV1 score have weaker effects (β_CD4,low CV1_ = 0.03, β_CD8,low CV1_ = 0.04, **Fig. 3D**). We observed that the CV1 interaction similarly superseded the discrete CD4+ interaction for 364/517 eQTLs with significant state dependence in both the discrete and continuous models, suggesting that the observed regulatory variation is more driven by a cell’s degree of cytotoxicity than by its lineage.

We hypothesized that multivariate modeling of multiple orthogonal CVs in an eQTL model could capture more granular, multifaceted states and their effects on gene regulation in individual cells. By sequentially adding CVs to a PME model and quantifying their cumulative significance, we observed the number of interacting eGenes reaches a maximum with 7 CVs in the model (2,237 interacting eGenes out of 6511, at LRT q<0.05, **Fig. 4A, Table S14**); adding more CVs does not substantially change the number of state-dependent eGenes (with 15 CVs: 2,221 interacting eGenes at q<0.05). CV interaction effects from the multivariate model were generally highly concordant with effects from univariate interaction models (r=0.87-0.97, **Fig. S9A, Table S15-20**), consistent with the independence of orthogonal CVs.

**Fig. 4.**
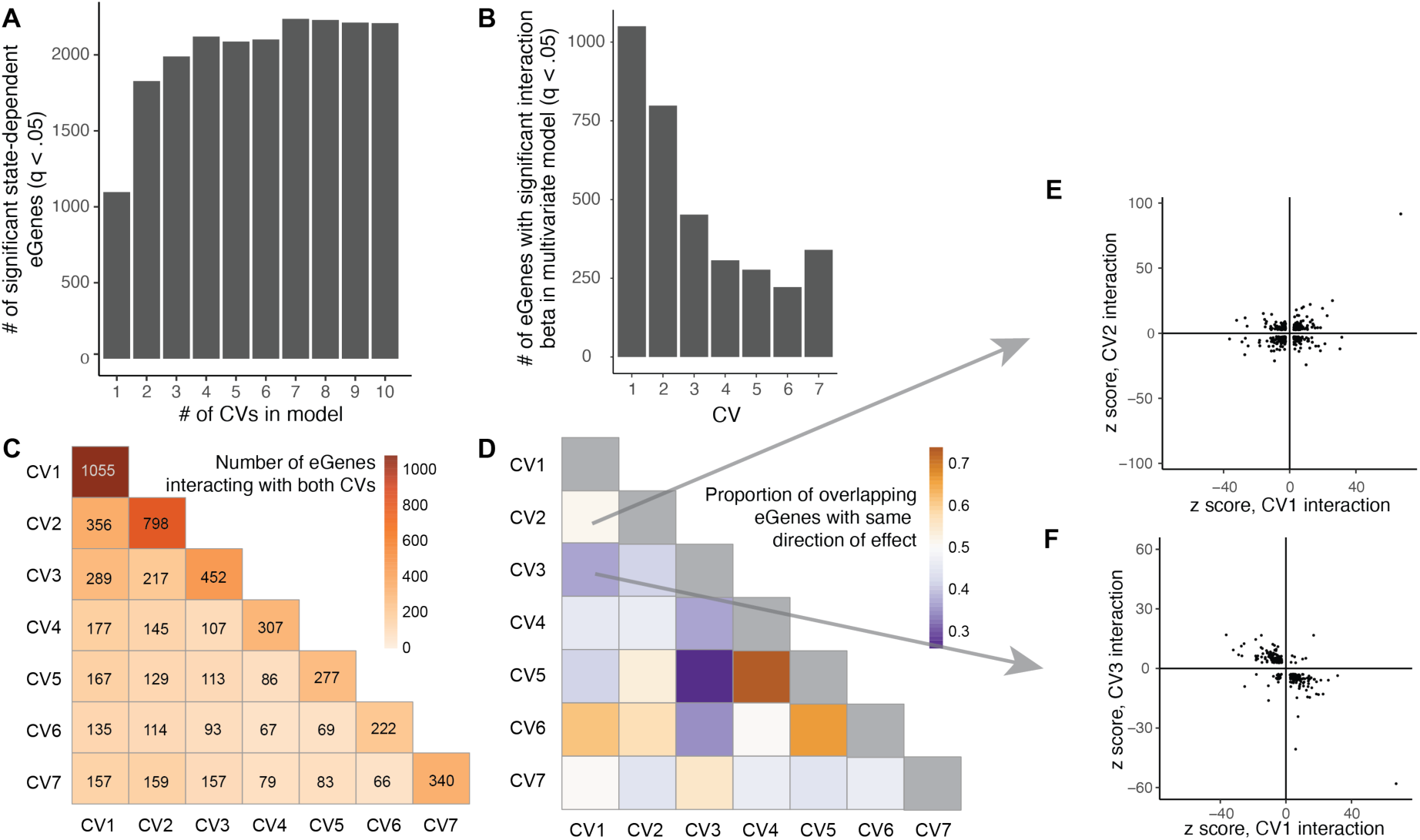
Cell-state-dependent eQTL interactions with continuous CVs. **(A)** Number of significant eGenes (LRT q < 0.05) detected by PME interaction models with increasing numbers of CVs. **(B)** Number of eGenes with significant interaction with each CV in a multivariate PME model with 7 CVs. **(C)** Heatmap of the number of eGenes with significant interactions with pairs of CVs in the multivariate model. Boxes along the diagonal reflect the total number of eGenes interacting with the corresponding CV. **(D)** Proportion of eGenes in **(C)** with the same direction of effect. **(E)** eGenes with significant interactions with either CV1 or CV2 plotted based on z scores with CV2 and CV1. **(F)** eGenes with significant interactions with either CV1 or CV3 plotted based on z scores with CV3 and CV1.

CV1 had the most interacting eGenes in both the univariate and multivariate (7 CV) models (**Fig. 4B, Fig. S9B**). Some eGenes significantly interacted with multiple cell states (**Fig. 4C**), and some pairs of states had related directions of effect in the multivariate model: for example, CV1 (cytotoxicity) and CV6 (T_H_1) tended to have the same direction of effect, while CV1 and CV3 (central) tended to have opposite directions (**Fig. 4D-F**). By clustering genes based on their interaction z scores (relative to the direction of the main effect) for the 7 CVs in the multivariate model, we defined eight broad clusters of genes based on distinct patterns of CV interactions that may reflect shared cell-state-dependent regulatory mechanisms (**Fig. S10, Table S21**).

### Individual cells have distinct eQTL effects

To estimate the eQTL effect for each gene at single-cell resolution, we can sum the products of interaction betas and corresponding CV scores for each individual cell (**Fig. 5A, Methods**). These CV scores capture the partial influence of each state that may be modulating regulatory activity. Adding this value to the baseline genotype beta estimates the total cell-level eQTL effect. These single-cell eQTL effects vary across cells and are independent of eGene expression.

**Fig. 5.**
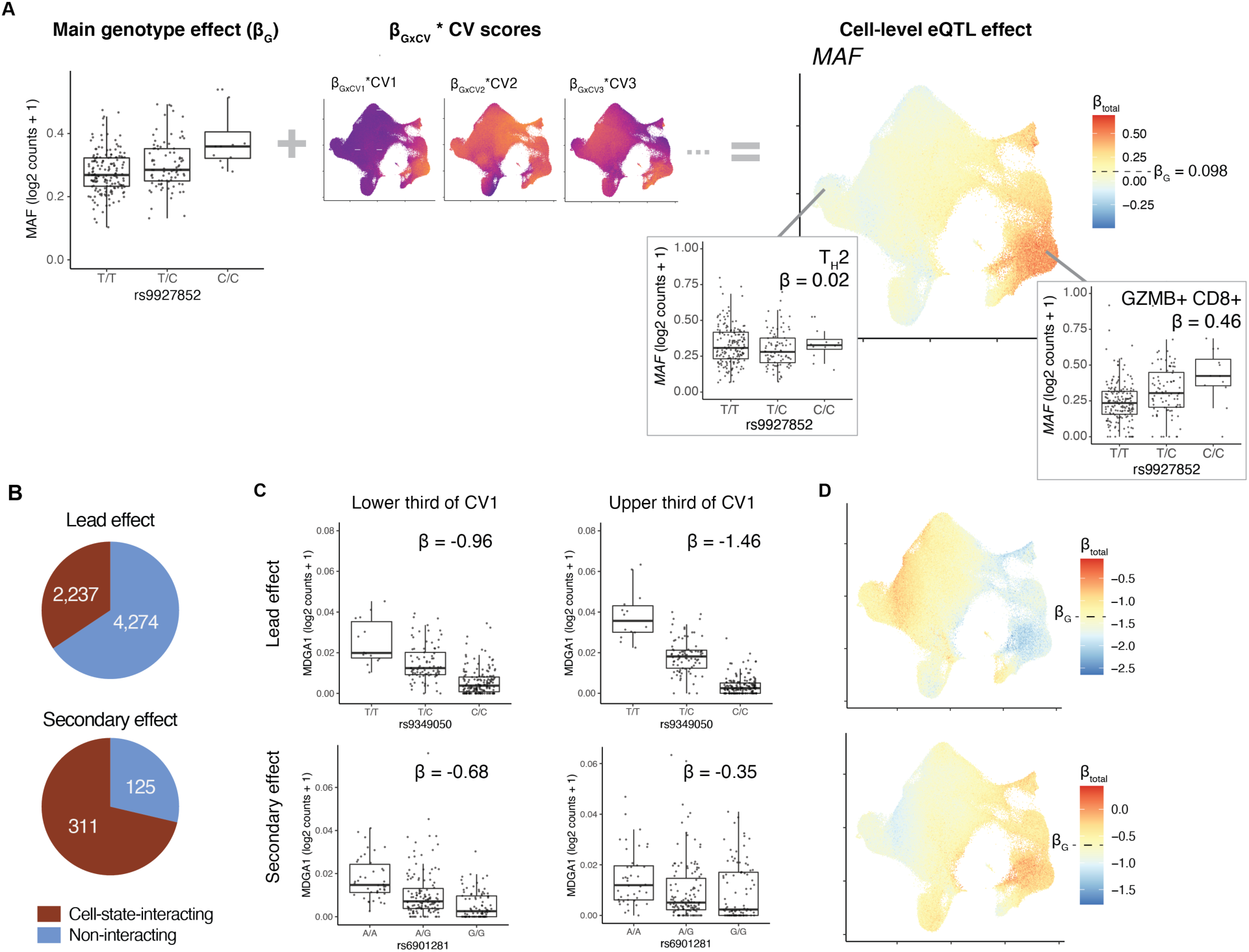
Single-cell dissection of eQTLs. **(A)** Schematic of calculating cell-level eQTL betas from main effect and CV interactions for the example of *MAF* and rs9927852. UMAP on the right shows total eQTL effect size at single-cell resolution, computed by summing main genotype effect (box plot) and individual CV effects (UMAPs). CV UMAPs depict each CV’s interaction beta multiplied cell-level CV scores scaled independently from lowest (purple) to highest (yellow). **(B)** Number of lead eQTL variants and independent secondary variants with significant cell-state interactions (red). (**C**) Lead (top) and secondary (bottom) eQTLs for *MDGA1* in cells with the lower third (left) and upper third of CV1 scores (right). Each point represents the average log2(UMI counts + 1) across all cells in the indicated CV1 score bin in a donor (n = 259), grouped by genotype. Box plots show median (horizontal bar), 25th and 75th percentiles (lower and upper bounds of the box, respectively) and 1.5 times the IQR (or minimum/maximum values if they fall within that range; end of whiskers). Beta values are the average β_total_ for all cells in the bin. (**D**) UMAP of total eQTL effect of lead (top) and secondary (bottom) variants for *MDGA1*. Each cell is colored by its β_total_, scaled to be centered on β_G_ with max (red) and min (blue) determined by the most extreme absolute β_total_ for that eQTL in any cell.

### Independent variants acting on the same eGene may have different state dependencies

Previous studies suggest that secondary eQTLs identified after conditioning on the lead effect are more likely to be cell-state-specific (*31*). We indeed observed a significantly larger fraction of secondary eQTLs with significant cell state interactions compared to lead variants (**Fig. 5B**, Fisher p = 6.98 x 10^-52^). Of the 436 secondary variants, 71% had cell-state specific effects compared to 34% of lead variants. 212 eGenes had at least two independent state-interacting effects. In some cases, eGenes’ lead and secondary variants may have contradictory interactions with the same CV. For example, the effect of *MDGA1*’s lead variant increases with CV1, while its secondary effect decreases (**Fig. 5C-D**). For *GNLY*, the effect of the lead variant increases with CV4, while its secondary effect decreases. Of the 64 eGenes with at least two independent effects interacting with CV1 (q < 0.05), 30 eGenes had different directions of CV1 interactions for their lead and secondary variants. Across all the CVs, 70 eGenes displayed this discordance with at least one state, demonstrating that cell states may not influence all the eQTLs for a gene in the same way **(Fig. S11).**

### State-dependent eQTLs colocalize with autoimmune-associated variants

Consistent with previous studies, we found that the memory T cell eQTLs that we defined with pseudobulk analysis were enriched for variants in LD with genome-wide significant loci associated with immune traits compared to genome-wide significant loci for all other traits in the GWAS catalog. For example, we observed relative enrichment for rheumatoid arthritis (RA; OR = 4.67, Fisher p = 2.25 x 10^-7^) and inflammatory bowel disease (OR = 4.84, Fisher p = 2.16 x 10^-11^, **Fig. S12A, Table S22**). We recapitulated previously described disease-associated eQTL variants, like rs1893592 (chr21_42434957_A_C), an eQTL for *UBASH3A* that is associated with RA (*32*).

We then assessed the cell-state dependence of disease-associated variants that overlapped eQTLs. Cell-state-interacting memory T cell eQTLs were enriched for overlap with GWAS variants compared to non-interacting eQTLs (OR = 1.31, Fisher p = 2.7 x 10^-4^), and state-dependent eQTL variants overlapped with at least one GWAS variant for 189/195 traits tested from the GWAS Catalog (*33*). State-interacting eQTLs were nominally enriched compared to non-interacting eQTLs for overlap with 14 individual traits, with associations that exceeded the null expectation (2,237/6,511 = 34%) for immune traits like RA (17 interacting, 7 non-interacting), type 1 diabetes (13 interacting, 7 non-interacting), and multiple sclerosis (24 interacting, 19 non-interacting) (ORs: 1.58-9.57; **Fig. S12B**, **Table S23**).

For example, the lead eQTL variant for *ORMDL3* (rs4065275) was in LD (r^2^ = 0.69 in 1KG PEL, r^2^ = 0.68 in 1KG EUR) with an RA GWAS variant (rs59716545) and also had a significant cell state interaction across CVs 1-7, driven by significant interactions with CVs 1 and 2 (*34*). The *ORMDL3* eQTL was strongest in the *GZMB*+ cytotoxic CD8+ T cells, intermediate in T_H_17s and other helper CD4+ states, and weaker in *RORC+* T_regs_ (**Fig. 6A**). On the other hand, the lead *IL18R1* eQTL variant (rs11123923, chr2_102351384_C_A)—in LD with inflammatory bowel disease GWAS variant rs1420098 (r^2^ = 1.00 in 1KG PEL and EUR)—was strongest in T_H_2s and T_H_17s with weaker effects in cytotoxic states (**Fig. 6B**) (*35*).

**Fig. 6.**
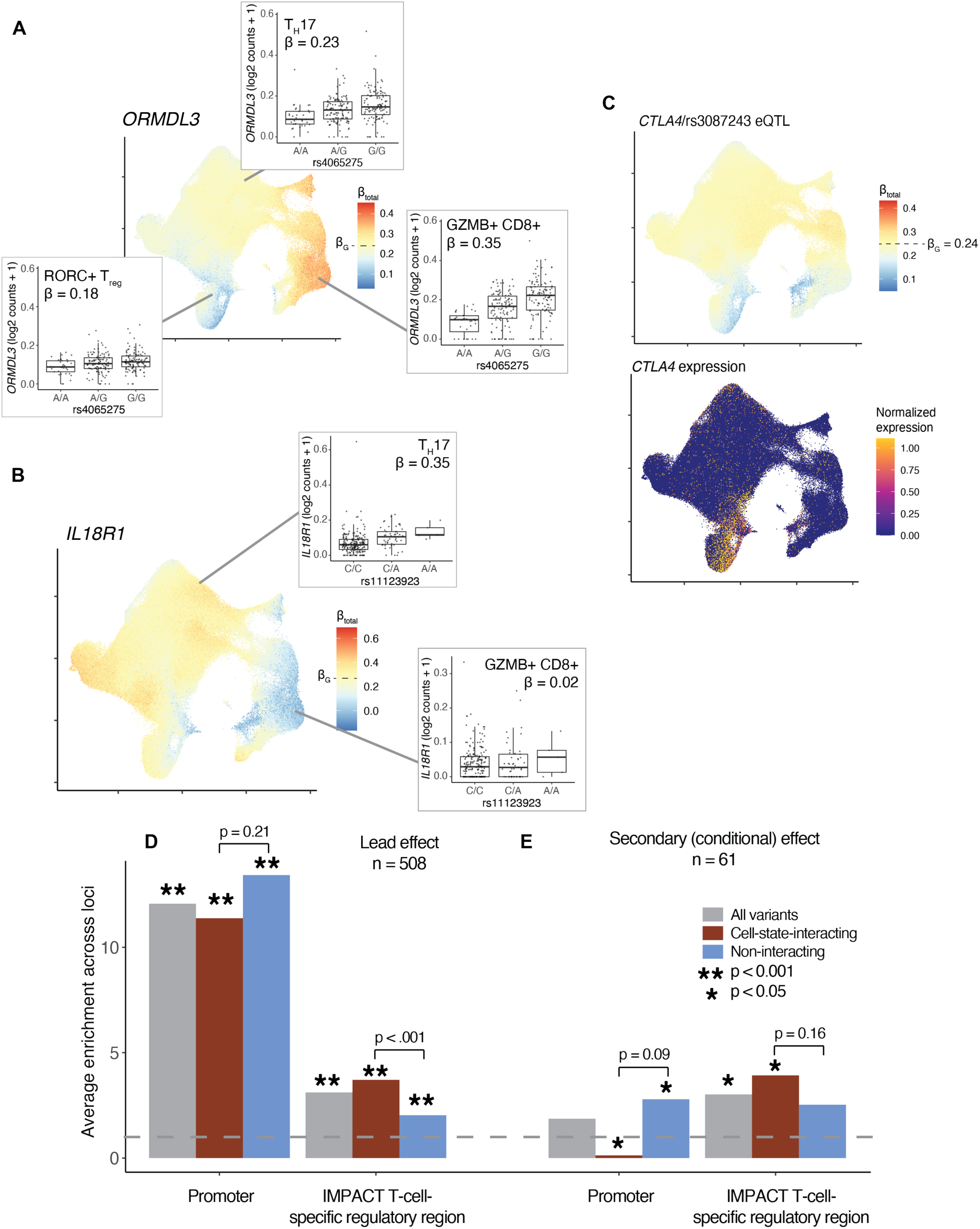
Cell-state dependent disease and regulatory impact of eQTLs. **(A)** UMAP of total effect size of RA-associated eQTL at rs4065275 for *ORMDL3* and **(B)**, IBD- associated eQTL at rs11123923 for *IL18R1* at single-cell resolution. **(C)** (top) UMAP of total effect size of RA-associated eQTL at rs3087243 for *CTLA4* at single-cell resolution. (bottom) Single-cell normalized expression of *CTLA4* gene scaled from lowest (dark blue) to highest (yellow). For all single-cell eQTL effect UMAPs, each cell is colored by its β_total_, scaled to be centered on β_G_ with max (red) and min (blue) determined by the most extreme absolute β_total_ for that eQTL in any cell. For all box plots, each point represents the average log2(UMI counts + 1) across all cells in the indicated cluster in a donor (n = 259), grouped by genotype. Box plots show median (horizontal bar), 25th and 75th percentiles (lower and upper bounds of the box, respectively) and 1.5 times the IQR (or minimum/maximum values if they fall within that range; end of whiskers). Beta values are the average β_total_ for all cells in the cluster. (**D),** Enrichment of eQTL lead effects or **(E)**, independent secondary (conditional) effects in promoter or T-cell-specific regulatory regions. Analysis was limited to loci where at at least one variant had PIP >= 0.5. The height of the gray bar corresponds to the average enrichment calculated across all loci containing a variant with PIP < 0.05, red bar corresponds to the subset with significant cell-state interaction (LRT q < 0.05 in model with 7 CVs), and the blue bar corresponds to the subset without significant cell-state interaction. Bars marked with an asterisk are have a one-sided permutation p value < 0.001. Each pair of interacting/non-interacting bars is labeled with a one-sided permutation p value for the difference (interacting minus non-interacting). The gray dotted line indicates enrichment statistic = 1.

GWAS variants did not always have stronger eQTL effects in states with higher overall expression. For example, the lead eQTL effect for *CTLA4* was mediated by rs3087243 (chr2_203874196_G_A), which is associated with RA (*36*). Although *CTLA4* expression is highest in a subset of Tregs, *RORC*+ Tregs, and activated CD4+ T cells, these cells had weaker eQTL effects (**Fig. 6C**). The eQTL effect was strongest in cytotoxic CD4+ T cells, a state with very low *CTLA4* expression. Our results suggest that disease processes may, in fact, emerge in unlikely states when pathogenic variants modulate low-level gene expression.

### State-dependent eQTLs are enriched in T cell regulatory regions

State-dependent eQTLs may be concentrated in particular regulatory regions, including promoters (whose effects are shared across states) or enhancers (which tend to have state-specific regulatory functions) (*37*). To test these regions for enrichment, we first defined promoters as the region within 2 kb of the transcription start site. We fine-mapped the eQTL effect at each locus with CAusal Variants Identification in Associated Regions (CAVIAR) based on summary statistics from the pseudobulk analysis (*38*). For loci where we were able to fine-map the lead effect to a single variant (n = 508, posterior inclusion probability [PIP] ≥ 0.5), we calculated a 12.07-fold enrichment of eQTL variants weighted by their PIPs in promoters (permutation p < 0.001, **Fig. 6D, Methods**). Cell-state-interacting and non-interacting eQTLs were both strongly enriched at 11.38- and 13.43-fold, respectively (p < 0.001, one-sided p for Δ_int-no int_ = 0.21), reflecting the regulatory importance of promoters regardless of state.

Next, we hypothesized that enhancers may also be enriched for cell-state specific eQTLs. Since there is uncertainty in the specific location of regulatory regions outside of the promoters, we defined cell-type-specific regulatory regions with Inference and Modeling of Phenotype-related ACtive Transcription (IMPACT), a logistic regression model trained on cell-type-specific transcription factor binding and epigenetic features (*39*). The IMPACT model for lineage-determining T-bet in CD4+ T_H_1 cells estimates the probability that the epigenetic landscape at any genomic position is favorable for transcription factor binding, where higher scores represent T-cell-specific regulatory regions. We only considered regions outside the previously defined promoters (TSS +/- 2kb) as T-cell-specific regulatory regions. We found that eQTL variants were enriched 3.12-fold (p < 0.001, **Fig. 6D**). Cell-state-interacting eQTLs were almost twice as enriched (3.71) as non-interacting eQTLs (2.03) in T-cell-specific regions (both p < 0.001, one-sided permutation p for Δ_int-no int_ < 0.001).

To more precisely identify causal variants based on effects shared across ancestries, we combined this dataset with European data from BLUEPRINT and conducted multi-ancestry fine-mapping of pseudobulk effects (*40*). We identified a single causal variant (PIP **≥** 0.5) explaining the lead effect for 1,247 eGenes also identified in the Peruvian analysis. As in the Peruvian analysis, these variants were enriched in promoters (13.94, p < 0.001) and T-cell-specific regulatory regions outside the promoter (2.94, p < 0.001), with greater enrichment of state-interacting eQTLs (3.74) in T-cell-specific regions compared to non-interacting eQTLs (2.06) (both p < 0.001, one-sided p for Δ_int-no int_ < 0.001; **Fig. S13**).

Previous studies have found that secondary eQTL variants are more likely to affect enhancers than promoters (*33*). Consistent with this, we found that relative to lead variants, secondary eQTL variants were less enriched in promoters (1.87, p = 0.12) and comparably enriched in T-cell-specific regulatory regions (3.02, p = 0.054) (**Fig. 6E**). However, only non-interacting secondary variants were enriched in promoters (2.79, p = 0.034), while cell-state-interacting secondary variants were significantly depleted (0.13, p = 0.018). There was no difference in enrichment in T-cell-specific regions between interacting (3.92, p = 0.008) and non-interacting variants (2.53, p = 0.056) (one-sided p for Δ_int-no int_ = 0.16). This suggests that secondary variants generally have more cell-type-specific regulatory roles, regardless of cell-state-dependence, but those found to be state-dependent in the PME model are especially depleted for shared effects.

## Discussion

Large single-cell datasets from genotyped cohorts—some with multiple single-cell data modalities—are becoming more common and make it possible to investigate how cell states shape the complex relationship between genetic variation, gene expression, and disease. In this study, we underscore the untapped potential of these data to reveal state-dependent regulatory heterogeneity when analyzed with traditional bulk methods and the urgent need to refocus eQTL analyses at single-cell resolution.

Recognizing growing evidence that clusters obscure the rich functional diversity of T cells and other dynamic cell types such as stem cells, stromal cells, and neurons, here we leveraged the granularity of single-cell data to better define state-dependent eQTLs (*41–43*). A single-cell Poisson model is computationally expensive—as noted by the few previous studies that have had mixed success with similar approaches—but its advantages over more common alternatives were most clear when we assessed state dependence: pseudobulk linear models cannot accommodate cell states defined at single-cell resolution, and a single-cell linear model was confounded by differential expression between states (*18, 44*). PME’s flexibility and robustness are important assets for effective state-dependent eQTL analysis.

Modeling continuous cell states in the PME model explained more overlooked variation, for example in rarer states like cytotoxic CD4+ T cells, which have been traditionally aggregated with other CD4+ T cells despite bearing more regulatory similarity to CD8+ T cells. This highlights the limitations of traditional discrete T cell states, which ignore the continuous ranges of T cell functions like activation, cytotoxicity, or helper lineages. However, continuous cell states can be difficult to interpret biologically, especially as more dimensions are considered jointly. We used multimodal CCA of gene expression and surface proteins for more robust definition and easier interpretation of these states. For other data modalities or cell types, alternative integration strategies may be effective (*45, 46*). Single-cell trajectories may reveal eQTLs varying along a unidimensional axis, like a perturbation or differentiation (*17*).

With these strategies, we identified state-dependent effects in a substantial proportion of eQTLs and reconstructed their joint effects in individual cells. Single-cell-resolution eQTL betas estimate the effect of a genetic variant on gene expression in any cell with a given cell-state profile, revealing that genetic variants can have different effects even in pairs of cells that share aspects of their states (for example, in the same cluster). We can not only identify conventional clusters in which disease-associated variants have different effects—such as CD8+ GZMB+ T cells for RA-associated rs4065275 near *ORMDL3*—but unbiasedly disentangle specific continuous states driving the overall eQTL that may transcend clusters, such as cytotoxicity for rs4065275 and *ORMDL3*.

These T cell states may be driven by distinct regulatory architectures, including transcription factors, epigenetic profiles, or chromatin accessibility patterns. State-dependent eQTLs may be in genomic positions that are only involved in regulatory activity in certain states—and the loci with independent eQTLs that have opposing state-dependent effects suggest that the exact position or nature of a variant determines these regulatory interactions. This study offers a starting point to design studies that further probe single-cell regulatory heterogeneity. For example, integrating single-cell ATAC-seq with RNA-seq in eQTL studies may offer insight into the overlap between these variants and state-specific accessible chromatin, or incorporating interactions between states or with abundance of a cell state to understand how the immune milieu shapes eQTL effects. Single-cell-resolution eQTLs can introduce new paradigms of how genes are regulated across diverse cell states.

## Materials and Methods

### Single-cell RNA-seq and genotype data and quality control (QC)

We previously published a dataset of memory T cells from a 259-donor subset of a Peruvian tuberculosis disease progression cohort (128 former cases, 131 former latently infected controls; GEO: GSE158769) along with detailed sample processing methods (*6, 42*). Briefly, we negatively isolated memory T cells with a modified Pan T cell magnetic-activated cell sorting (MACS^R^) kit with anti-CD45RA biotin and followed an optimized version of the CITE-seq protocol with TotalSeq^TM^-A (BioLegend) oligonucleotide-labeled antibodies for a panel of 31 surface proteins (*43*). We pooled cells into batches of six donors for 10x Genomics library preparation and sequenced on an Illumina HiSeq X. Reads were aligned to GRCh38 with Cell Ranger. After demultiplexing donors with Demuxlet, we removed cells labeled as doublets, with < 500 genes expressed or > 20% of unique molecular identifiers (UMIs) from mitochondrial genes, from samples whose genotypes did not match genotypes called from single-cell data, or lacking surface markers of memory T cells (CD3 and CD45RO) (*44*).

A superset of 4,002 donors was genotyped in a separate genetic study on a custom Affymetrix array (LIMAArray) based on whole exome sequencing from 116 individuals with active TB from the same Peruvian cohort (dbGaP: phs002025). The design of this array has been described previously (*45*). We removed variants that were significantly associated with batch (p < 1 x 10^-5^), duplicated, or had low call rate, significant differences in the missingness rate between cases and controls (> 10^-5^), or Hardy-Weinberg p value < 10^-5^ in controls.

We mapped variants to GRCh37/b37 and used SHAPEIT2 to pre-phase genotypes and IMPUTE2 to impute genotypes with 1000 Genomes Project Phase 3 as the reference panel (*46, 47*). After removing SNPs with an INFO score < .9, minor allele frequency < 0.05, or deletions, the remaining variants were converted to GRCh38 with liftOver.

After single-cell and genotype QC, we used 500,089 cells from 259 donors and 5,460,354 variants for eQTL analysis.

### Single-cell data processing

mRNA and protein data were processed separately, as described (*6*). Briefly, we normalized the UMIs for each gene in each cell to log(counts per 10,000) and used centered-log-ratio normalization for each protein within each cell. Normalized mRNA and protein expression were scaled so that each feature had mean = 0, variance = 1 across all cells. After selecting the top 1,000 variable genes per donor and removing the mouse immunoglobulin G protein (control), we conducted principal component analysis of each modality with the irlba R package and corrected the top 20 PCs of each modality for donor and library preparation batch effects with Harmony (*48*).

### Pseudobulk eQTL analysis

To make pseudobulk expression profiles for 259 donors, we removed cells from 12 technical replicate samples and summed the UMI counts for each gene across all cells from each donor, producing one aggregated expression value for each gene in each donor. For CD4+ and CD8+ pseudobulk analysis, we constructed pseudobulk expression profiles for each donor in each compartment by *in-silico* gating cells that were CD4+CD8- or CD8+CD4-, respectively, based on their normalized surface protein expression measured in CITE-seq. Gates were defined through visual inspection. For T_reg_ pseudobulk analysis, we used previously defined cluster annotations to construct a pseudobulk expression profile for the cells in clusters C-5 (*RORC+* T_reg_) and C-9 (T_reg_) (*6*).

Genes were removed if expressed in fewer than half the donors (pseudobulk counts > 0 in ≤ 129 donors). For the remaining 15,789 genes, we normalized the pseudobulk profiles to log_2_(counts per million + 1) and applied inverse normal transformation. Then, we used probabilistic estimation of expression residuals (PEER) implemented in R to regress out age, sex, five genotype PCs, and 45 PEER factors (*49*).

We then conducted a whole-genome eQTL analysis for all 22 autosomal chromosomes. For each gene, we associated its residual expression after PEER normalization with the dosage at each SNP within 1 MB of the transcription start site. These models were implemented in FastQTL (default settings) (*50*). To ensure robustness, we used FastQTL’s beta approximation to compute a permutation p value from 1,000 permutations. To correct for multiple hypothesis testing, we calculated q values for the lead SNP for each gene, and identified eGenes with significant eQTL variants with q < 0.05 (*51*).

For conditional pseudobulk analysis, we iteratively regressed each eGene’s PEER-normalized residuals on the dosage of its lead SNP and used the subsequent values for FastQTL analysis. We repeated this twice.

### Comparison to the BLUEPRINT project

To validate this model’s ability to detect previously characterized eQTLs, we used the naive CD4+ T cell eQTLs reported by the BLUEPRINT project (*15*). We selected eGenes that were significant in our dataset (n = 6,511) and identified the subset of eGene/lead variant pairs also measured by BLUEPRINT (n = 3,249) and significant in both datasets at q < 0.05 (n = 2,056). We compared the direction of effect for these variants between the two datasets.

### Continuous cell state definition and annotation

For multimodal dimensionality reduction, we used canonical correlation analysis, as implemented in the cc function from the CCA R package (*52*). We ran CCA on scaled mRNA expression for the most variable genes (excluding T cell receptor genes) and scaled protein expression for all 30 memory T cell proteins, and computed cells’ scores on each canonical variate (CV) based on the weight of each gene on each CV. We then corrected scores on the top 20 CVs for donor and batch effects. For visualization, we projected this embedding into a two-dimensional Uniform Manifold Approximation Projection (UMAP) with the umap function from the uwot R package (*53*).

To annotate each CV based on its biological correlates, we first measured the Pearson correlation coefficient between cells’ CV scores and the normalized expression of each surface protein marker we measured. We measured correlations between CV scores and the normalized expression of genes encoding lineage-defining genes.

We also conducted gene set enrichment analysis. First, we measured the correlation of each CV prior to batch correction with the expression of each gene used as input for CCA. These correlations defined the ranked gene list for each CV. Then, we measured the enrichment of each immunologic gene set (C7, only those annotated as “UP”) in MSigDB and a published “innateness” gene list in each CV’s ranked gene list with the fgsea function in the fgsea R package (*54, 55*). We corrected for multiple hypothesis testing with a Bonferroni p value threshold adjusted for 2,360 gene sets tested (0.5/2,360 = 2.12 x 10-5).

To compare CV scores between and within clusters, we used the cluster annotations defined in the previous study and computed the average score on each CV for cells from each cluster. We also randomly selected two cells each from clusters C-3 and C-15 to compare to each other and the cluster.

### Single-cell eQTL modeling

We modeled single-cell eQTLs with a Poisson model of each gene’s UMI counts as a function of genotype at the eQTL variant and other donor- and cell-level covariates, for each gene:

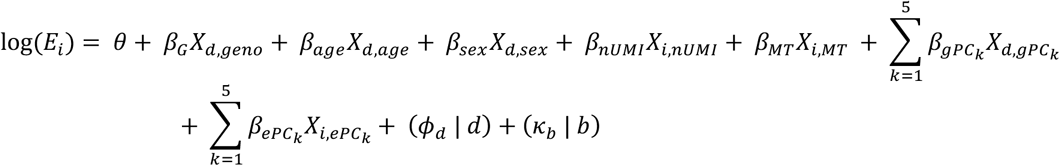

where *E* is the expression of the gene in cell *i*, θ is an intercept, and all other βs represent fixed effects as indicated (nUMI = number of UMI, MT = proportion of mitochondrial UMIs, gPC= genotype PC, ePC=single-cell mRNA expression PC) for covariates in cell *i*, donor *d*, or batch *b*. Donor and batch are modeled as random effect intercepts.

To test interactions with cell state, we added a fixed effect for cell state and a cell state x genotype interaction term:

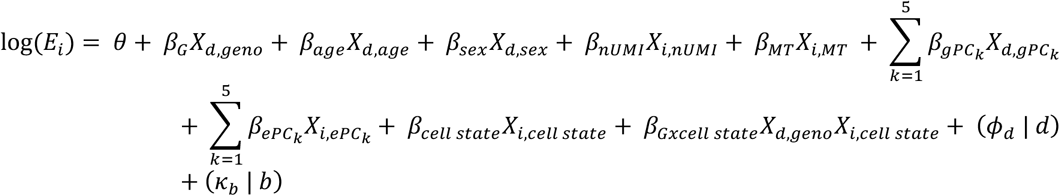

When testing whether the discrete state (CD4+) or the continuous state (CV1) captured more variance, we included cell state and cell state interaction terms for both and removed each interaction term to create the corresponding null model for the likelihood ratio test (described below).

To test interactions with former TB progression status, we used the same model but with a fixed effect and genotype-interaction term for TB status.

To test interactions with multiple state-defining covariates (e.g., multiple CVs), we included additive fixed effect and interaction terms for each CV:

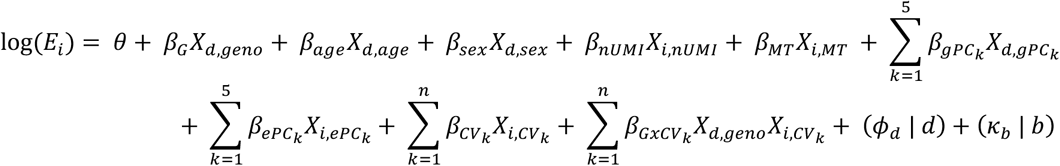

To test eQTLs within discrete cell states (e.g., CD4+), we subsetted the full dataset to cells in the state of interest (using gates or clusters). Then, we ran the Poisson single-cell model without any cell state terms.

We fit all single-cell Poisson mixed models with the glmer function in the lme4 R package, with family=“poisson”, nAGQ=1, and control=glmerControl(optimizer = “nloptwrap”) (*56*). To determine the significance of this model, we used a likelihood ratio test comparing the models with and without the genotype term (for the memory T cell analysis) or the cell state interaction term(s) (for the cell-state-specific analyses) and calculated a p value for the test statistic against the Chi-squared distribution with one degree of freedom. We corrected for multiple hypothesis testing by calculating q values across all tested eQTLs.

For comparison, we also used a single-cell linear mixed effects (LME) model to test eQTLs across all memory T cells and for state dependence with CD4+ state or continuous CV1. The model included the same covariates as the Poisson models. For example, across all memory T cells:

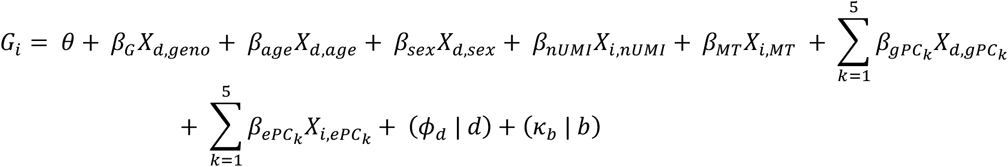

where *G* is the log2(counts per 10K) normalized expression of the gene in cell *i*. We fit all single-cell linear mixed models with the lmer function in the lme4 R package, with REML = F (*56*) and determined the significance of the model as described for Poisson models.

### Type 1 error estimation

To estimate the false positive rate for the single-cell PME model of memory T cell eQTLs, we permuted genotype across donors and ran the PME model for each gene (n = 6,511). Then, we used a likelihood ratio test to compute a p value for each gene under the permutation and measured the proportion of genes with a p value < alpha = 0.05. To estimate the false positive rate for the single-cell PME model of cell state-dependent eQTLs, we used the same permutation approach but permuted cell state across cells to preserve the main genotype effect.

### Simulating differential expression

We selected genes with non-significant cell state and cell-state interaction terms in the PME model with CD4+ as the cell state of interest. To test robustness of the single-cell models to differential expression, we uniformly reduced the expression level of each gene in each cell to 50%, 20%, and 10% of baseline expression. For PME, we did this in log_2_-space, converted back to counts, and rounded to the nearest whole number. For LME, we reduced expression in log2(counts per 10,000) space. Then, we ran the single-cell Poisson model with cell-state interaction (cell state = CD4+ cells).

### Clustering eQTLs and cells

We stringently selected cell-state-dependent eGenes (LRT p value from modeling CVs1-7 < 0.05/6511 = 7.7 x 10^-6^. For each eGene, we extracted the z score for each cell state interaction term from the Poisson model and multiplied them by the sign of the main genotype beta to standardize directions of effect, i.e., positive value means interaction amplifies baseline genotype effect, negative value means interaction dampens effect. Using Seurat, we built a shared nearest neighbor graph and used Louvain clustering with n.start = 20, n.iter = 20, and resolution = 1.5 to define eight clusters of eGenes (*57*).

We explored the potential biological significance of the clusters by measuring the enrichment of MSigDB Hallmark and Gene Ontology gene sets. For each gene set, we used a Fisher’s exact test to compare the proportion of eGenes in the cluster that overlapped with the gene set versus the proportion in other clusters that overlapped with the gene set. We assessed significance with a Bonferroni threshold of 0.05/14,765 gene sets tested = 3.4×10^-6^.

### Cumulative eQTL interaction effect

The effect of each eQTL in each cell is the cumulation of the main genotype effect and all of the genotype x CV interactions. We calculated the overall effect for each eQTL in each cell by summing the genotype beta and the products of each CV score and the corresponding beta:

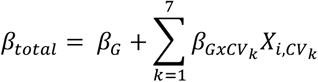

For interpretation in some analyses, we defined cluster-level betas by averaging cell-level betas for all cells assigned to that cluster in the previous study.

### GWAS variant enrichment

We downloaded the GWAS Catalog as of July 30, 2020, restricted to GWAS in European populations, and identified variants associated with each of 194 traits at p < 5×10^-8^ and pruned with plink to remove variants with LD r^2^ > 0.2 (*29, 58*). We also constructed a set of background variants by pooling variants associated with any trait at p < 5×10^-8^ and pruning with plink to remove variants with LD r^2^ > 0.2.

For each memory-T cell eQTL, we identified all other variants with LD r^2^ ≥ 0.5 in both the 1000 Genomes Peruvians in Lima, Peru (PEL) and European (EUR) populations. We then matched these variants with variants from the GWAS Catalog for 194 traits and the background variant set. We calculated all enrichments with a two-sided Fisher test. To calculate memory-T-cell eQTL enrichments for specific traits, we compared the proportion of eQTL-colocalizing GWAS variants for each trait with the proportion of eQTL-colocalizing background variants. To calculate the enrichment of GWAS variants colocalizing with state-dependent eQTLs, we compared the proportion of eQTLs with significant cell-state interaction (LRT q < 0.05 from the model with 7 CVs) that colocalize with the background variant set compared to the proportion of eQTLs without significant cell-state interaction that colocalize.

### Fine-mapping memory-T cell eQTLs

For each locus (eGene), we used CAusal Variants Identification in Associated Regions (CAVIAR) software allowing only a single causal variant in each locus (-c 1) to estimate the probability that each variant in a +/- 250kb window around the transcription start site (TSS) is causal (*34*). We ran CAVIAR on pseudobulk eQTL z scores for these variants and pairwise Pearson correlation coefficients between the variants (calculated with plink version v1.9b) (*58*).

For joint multi-ancestry analysis, we first lifted the BLUEPRINT dataset to GRCh38 with liftOver and filtered at a minor allele frequency threshold of 0.05. We then merged the TBRU and BLUEPRINT datasets matching on chromosome, position, reference, and alternate alleles and performed eQTL analysis on the joint datasets as described above. We fine-mapped each locus with CAVIAR using z scores from the joint analysis, as described in the Peruvian dataset.

### Enrichment of eQTLs in regulatory regions

We defined promoters as the region +/- 2kb from the transcription start site of each of the 6,511 significant eGenes based on the Cell Ranger 3.1.0 GTF. This annotation was binary.

We defined cell-state-specific regulatory regions with a probabilistic annotation of the genome by Inference and Modeling of Phenotype-related ACtive Transcription (IMPACT) (*35*). First, we collected public T-bet (*TBX21*) ChIP-seq data for in CD4+ T_H_1 cells from NCBI as a gold standard for CD4+ T_H_1 regulatory elements (*59*). We also previously aggregated 5,345 public epigenetic features from NCBI, ENCODE, and Roadmap spanning all possible cell types (*60*). Then, we used IMPACT’s logistic regression model to distinguish 1,000 T-bet bound sequence motifs from 10,000 unbound T-bet sequence motifs genome-wide based on epigenetic feature characterization. We used HOMER [v.4.8.3] to identify T-bet sequence motif matches as previously done (*35, 61*). We then characterized every nucleotide genome-wide using the same set of epigenetic features and estimated the probability (between 0 and 1) of a regulatory element important to the cell type.

For the multi-ancestry analyses, we restricted loci to those that were also significant eQTLs in the Peruvian-only eQTL analysis. We computed enrichments in each locus containing a variant with posterior inclusion probability ≥ 0.5 and averaged across loci. To compute enrichments for the binary promoter annotation, we determined whether each variant in the locus overlapped with a promoter region (X = 0 if no overlap, X = 1 if overlap). Then, we calculated the enrichment across n variants in the locus as:

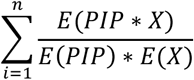

To compute enrichments for the probabilistic IMPACT T cell regulatory region annotations, we used the same strategy but X = the IMPACT score between 0 and 1 for each variant. We determined the significance of each enrichment by comparing the true enrichment score to a null distribution constructed by permuting the PIPs across variants in each locus 1,000 times and calculating an enrichment score. We determined the significance of the difference between the enrichments in interacting vs. non-interacting eGenes by comparing the true difference to a null distribution constructed by permuting interacting vs. non-interacting labels across eGenes and calculating an enrichment score. Both p values were computed with a one-sided comparison.

---

**Fig. S1.**
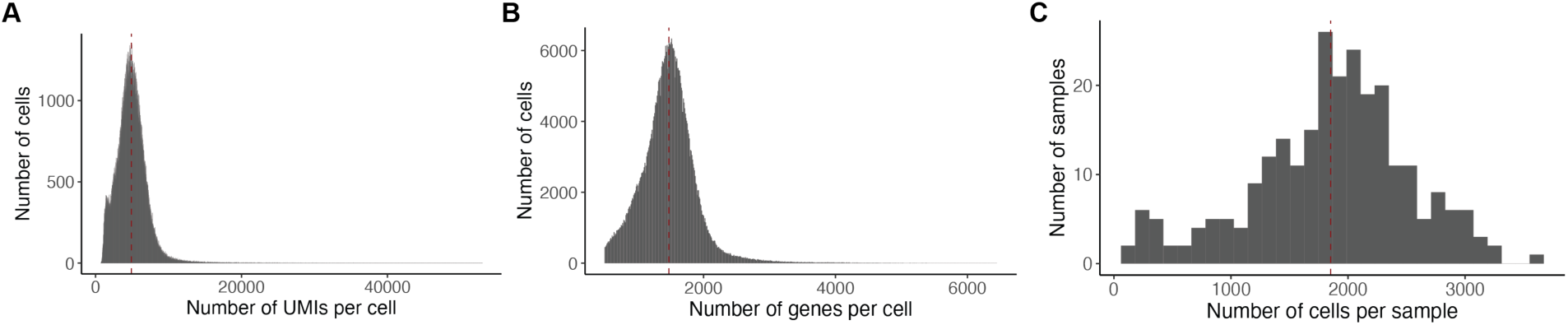
Single-cell quality control. **(A)** Histogram of 500,089 cells by number of UMIs and **(B)** number of genes. **(C)** Histogram of 259 samples by number of cells.

**Fig. S2.**
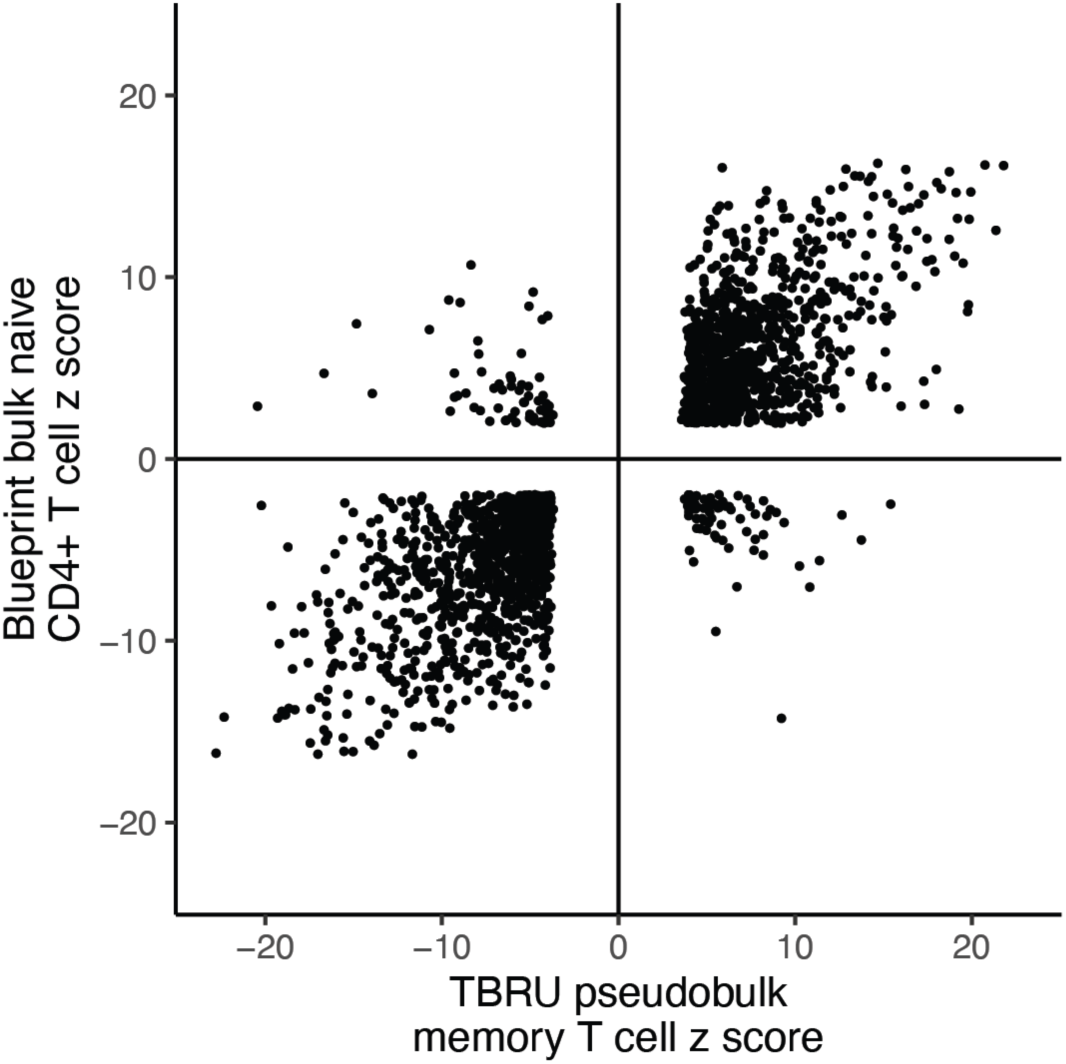
Concordance of BLUEPRINT and Peruvian (pseudo)bulk eQTLs. Each point represents an eGene/Peruvian lead variant pair significant in both data sets (q < 0.05), plotted based on the z score of the beta in BLUEPRINT vs. Peruvian data set.

**Fig. S3.**
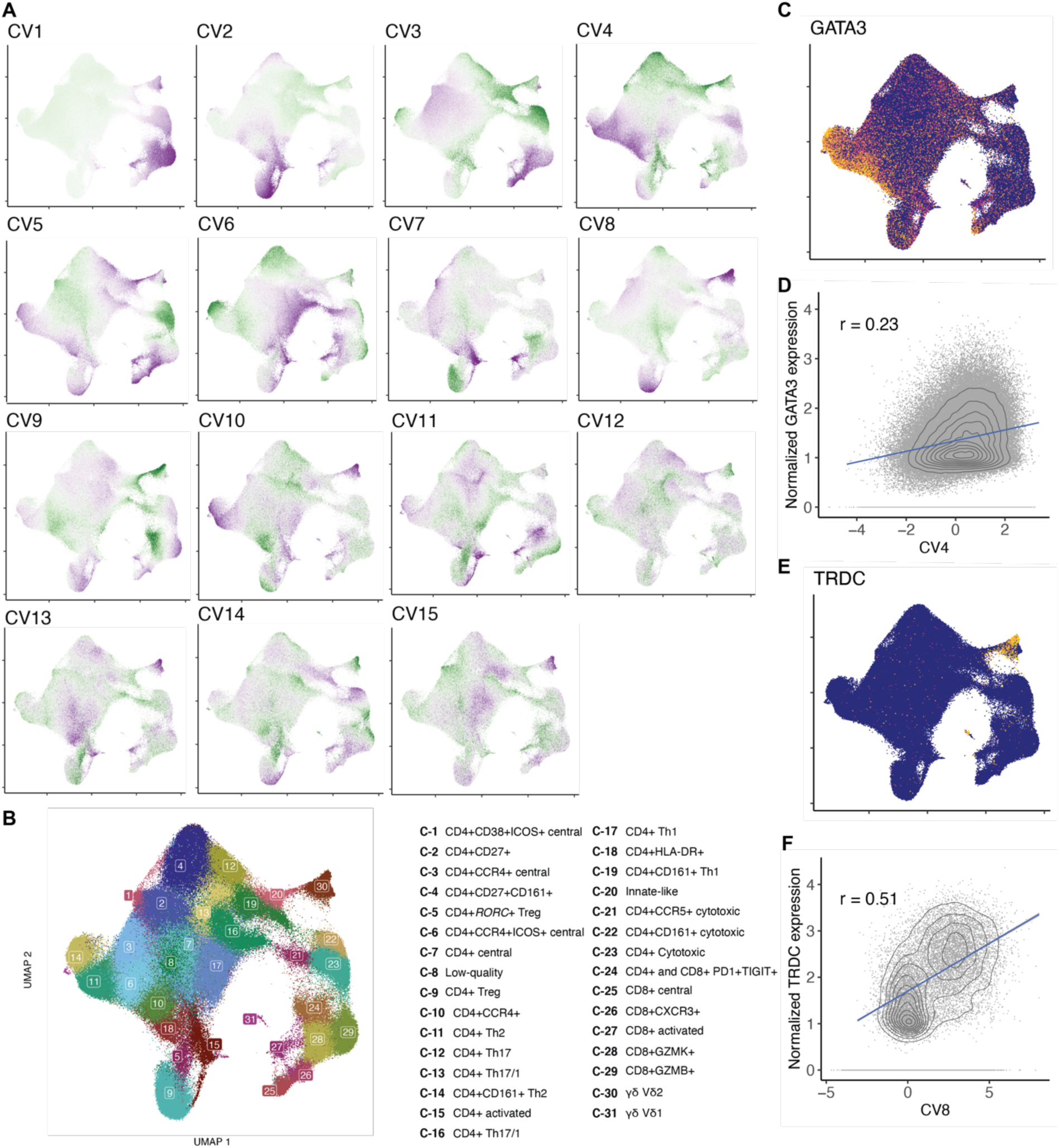
Continuous canonical variates (CVs) capture single-cell heterogeneity. **(A)** UMAP of memory T cells (n = 500,089) colored by score along each of the top 15 CVs from CCA-based integration of mRNA and protein expression in Nathan, et al. **(B)** UMAP of 31 memory T cell clusters defined with Louvain clustering in Nathan, et al. **(C)** and **(E)** UMAP of memory T cells colored by normalized expression (log2(UMI counts per 10,000)) for *GATA3* and *TRDC*. **(D)** and **(F)** Cells plotted based on normalized expression of *GATA3* and CV4 score or *TRDC* and CV8 score. For cells with non-zero expression, blue line represents linear best-fit line, contours represent density of cells, and r is the Pearson correlation coefficient.

**Fig. S4.**
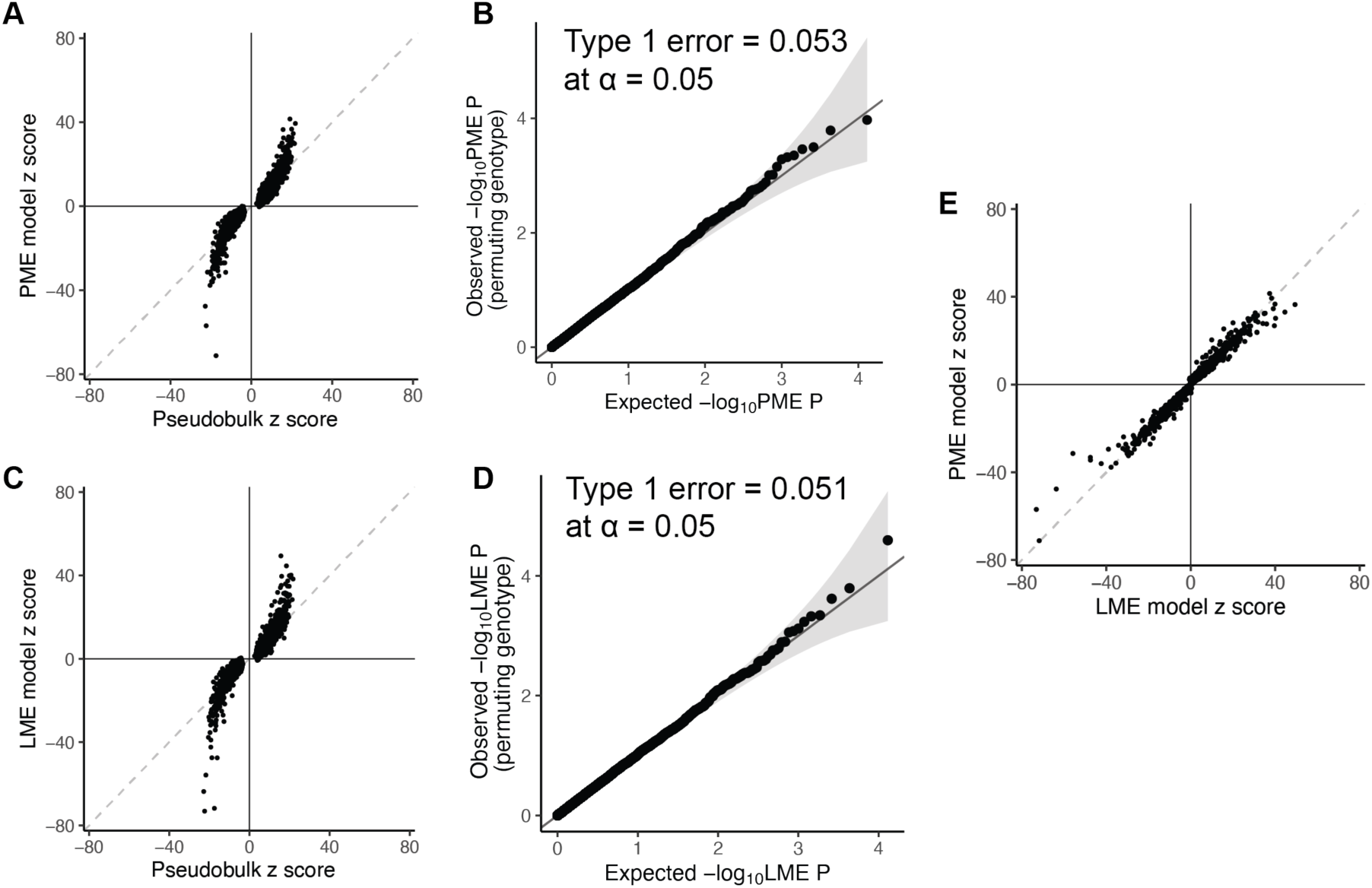
Comparing PME and LME models for single-cell eQTL analysis. **(A)** and **(C)** Pseudobulk-significant eGenes (n = 6,511) plotted based on z score from **(A)** PME or **(C)** LME single-cell model and pseudobulk model. Dashed line represents the identity line. **(B)** and **(D)** Quantile-quantile plot of pseudobulk-significant eGenes plotted based on -log10 p value of genotype beta from **(B)** PME or **(D)** LME model with permuted genotypes and uniformly distributed quantiles. Type 1 error is calculated as the proportion of eGenes with p < 0.05 with permuted genotypes. **(E)** Pseudobulk-significant eGenes plotted based on z score from PME and LME single-cell model

**Fig. S5.**
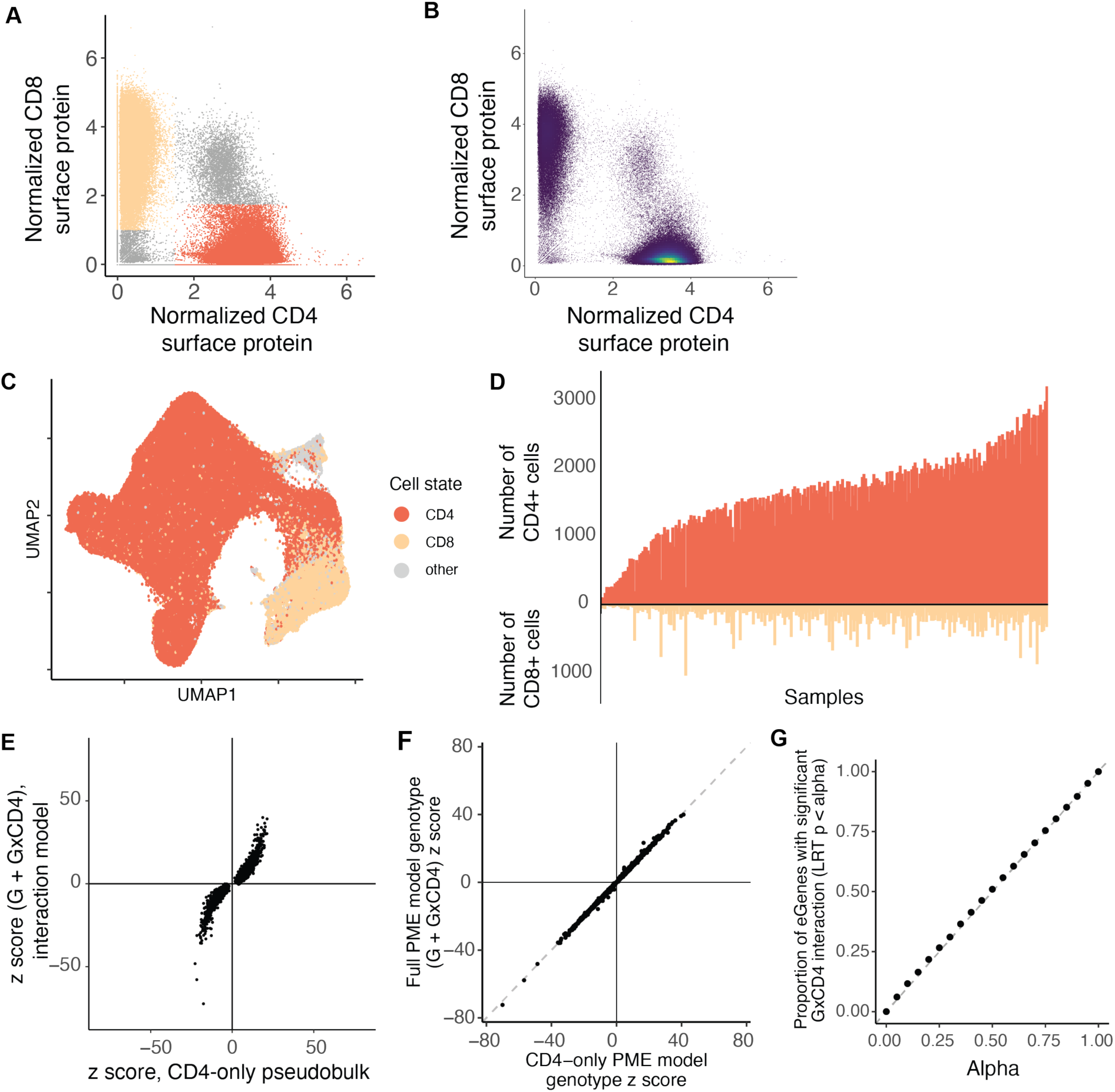
Modeling eQTL interactions with discrete CD4+ and CD8+ cell states. **(A)** Cells plotted by CLR-normalized CD8 and CD4 protein expression. Orange cells are CD4+CD8- and beige cells are CD4-CD8+ and in **(B)** cells are colored by density. **(C)** UMAP of 500,089 memory T cells colored by CD4+CD8- (orange) and CD4-CD8+ (beige). **(D)** Bar graph of number of CD4+CD8- cells (top) and CD4-CD8+ cells (bottom) per sample (n = 259). **(E)** and **(F)** Dot plot of each memory-T-cell eGene based on z score for total beta (β_G_+β_GxCD4_) in PME interaction model of all cells and z score for β_G_ from **(E)** pseudobulk model or **(F)** PME model of only CD4+ cells. **(G)** Proportion of eGenes with significant β_GxCD4_ under genotype permutation. Each dot represents the proportion significant at the given alpha threshold.

**Fig. S6.**
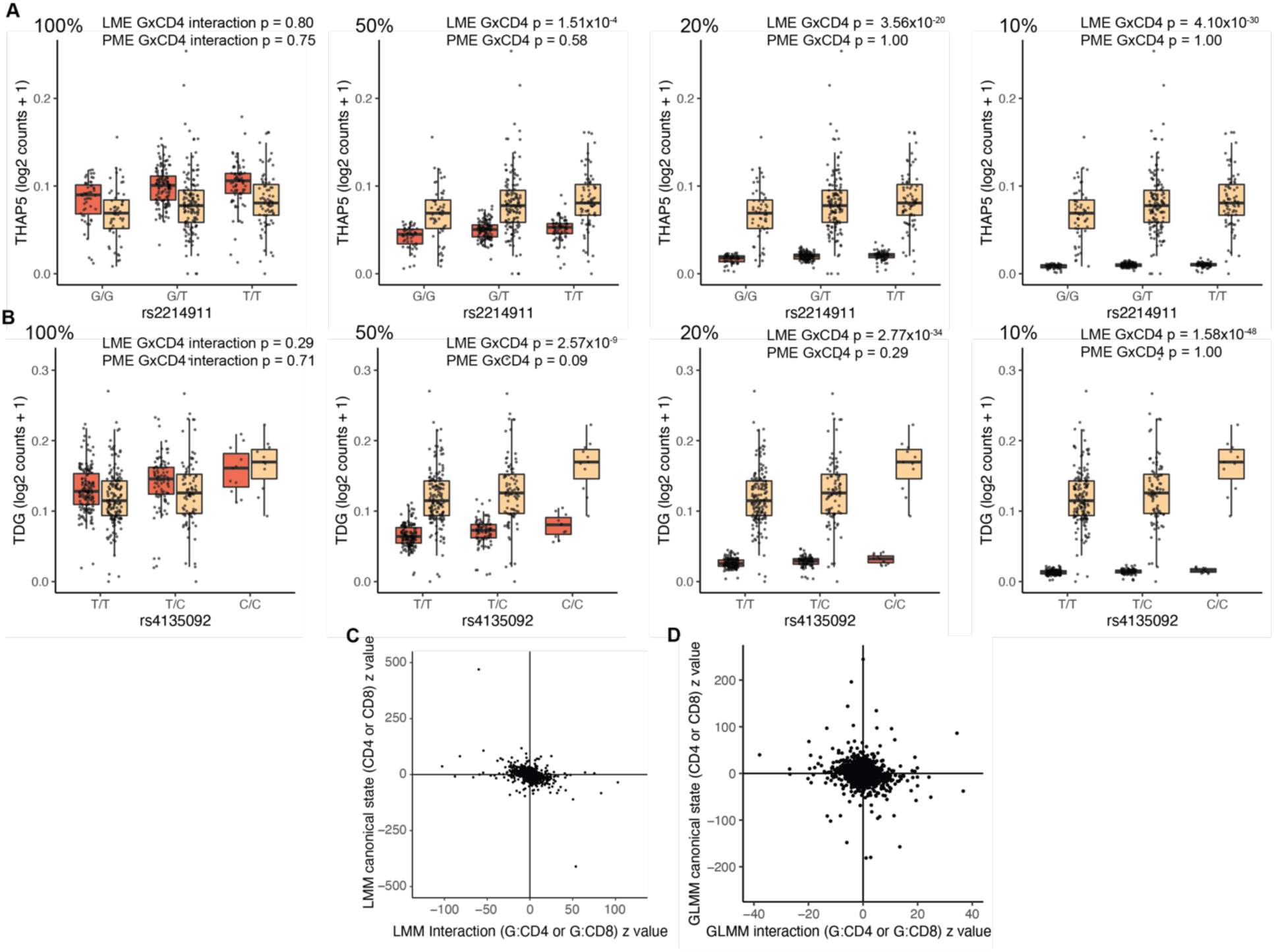
Assessing robustness of single-cell models to differential expression. (**A)** rs2214911 eQTL for *THAP5* and **(B)** rs4135092 eQTL for *TDG* in CD4+ (orange) and CD8+ (beige) cells. Box plots show the eQTL effects as per-cell gene expression decreased from 100% to 50, 20, and 10 percent in CD4+ cells (left to right). Each point represents the average log2(UMI counts + 1) across all cells in the indicated subset of cells in a donor (n = 259), grouped by genotype. Box plots show median (horizontal bar), 25th and 75th percentiles (lower and upper bounds of the box, respectively) and 1.5 times the IQR (or minimum/maximum values if they fall within that range; end of whiskers). **(C)** and **(D)** Dot plot of memory-T-cell eGenes (n = 6,511) based on z score for cell state beta (β_CD4_ or β_CD8_) and z score for cell state interaction beta (β_GxCD4_ or β_GxCD8_) in **(C)** LME and **(D)** PME models.

**Fig. S7.**
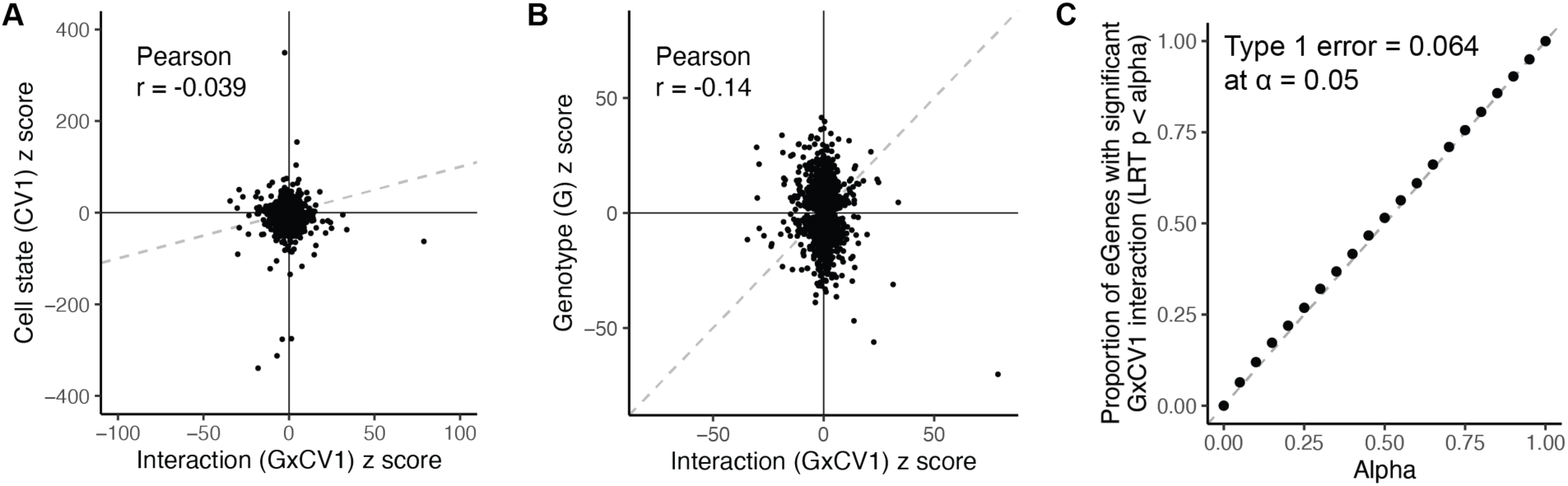
Calibration of the PME interaction model. Dot plot of memory-T-cell eGenes (n = 6,511) based on z score for **(A)** cell state beta (β_CV1_) or **(B)** genotype (β_G_) and z score for cell state interaction beta (β_GxCV1_) in PME model. **(C)** Proportion of eGenes with significant β_GxCV1_ under genotype permutation. Each dot represents the proportion significant at the given alpha threshold.

**Fig. S8.**
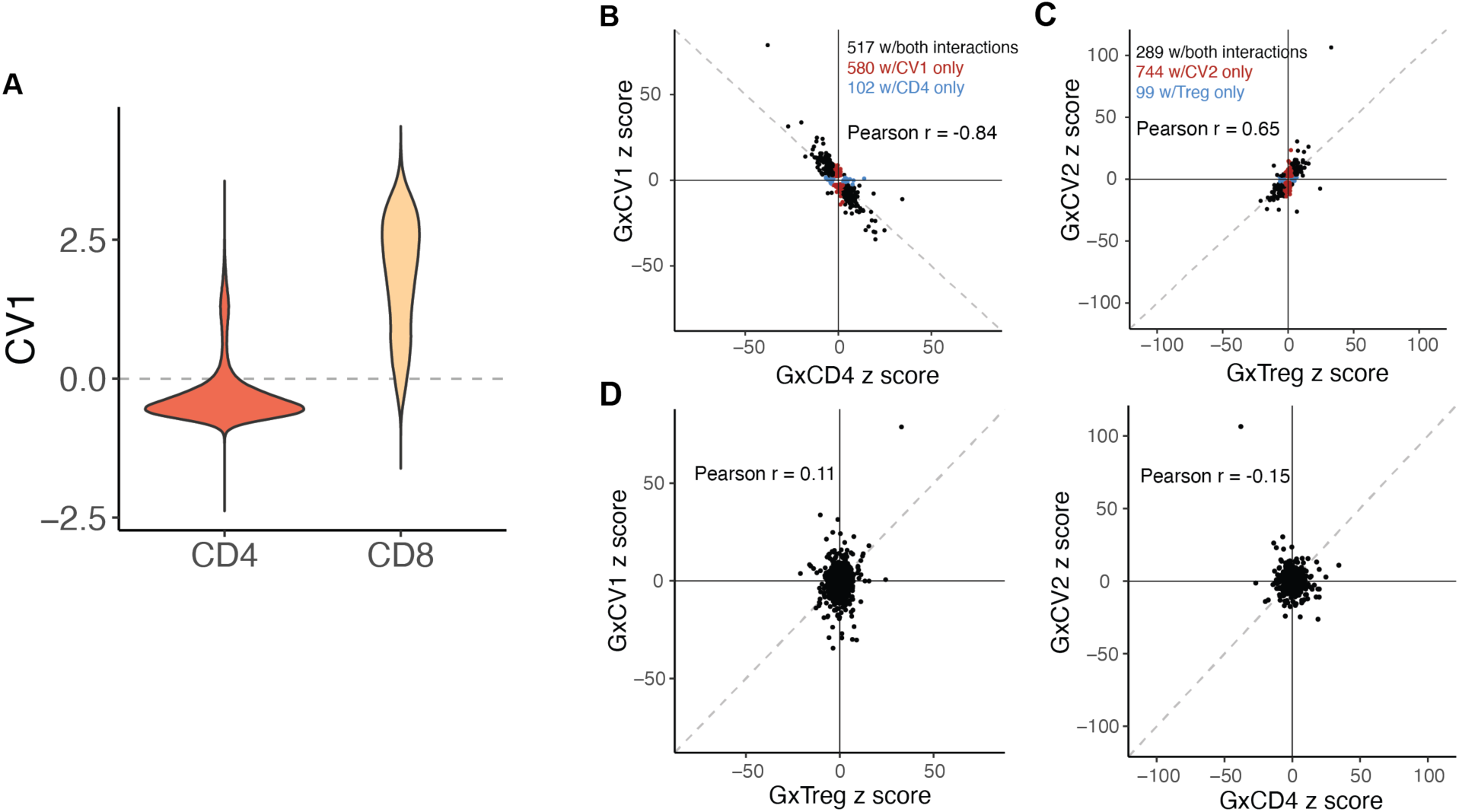
Concordance between eQTL interactions with continuous and discrete states. **(A)** Distribution of CV1 scores for cells in CD4+ and CD8+ gates. Dashed line represents CV1 = 0. **(B)**-**(E)**, Dot plots of eGenes’ z scores of genotype interactions with **(B)** CV1 and CD4+, **(C)** CV2 and Treg, **(D)** CV1 and Treg, or CV2 and CD4+. Dashed line represents the identity line. Only eGenes with significant interaction (LRT q < 0.05) are plotted in **(B)** and **(C)**. Black dots represent eGenes significantly interacting with both continuous and discrete states, red dots are only significantly interacting with continuous state, and blue dots are only significantly interacting with discrete state. r is calculated as the Pearson correlation coefficient.

**Fig. S9.**
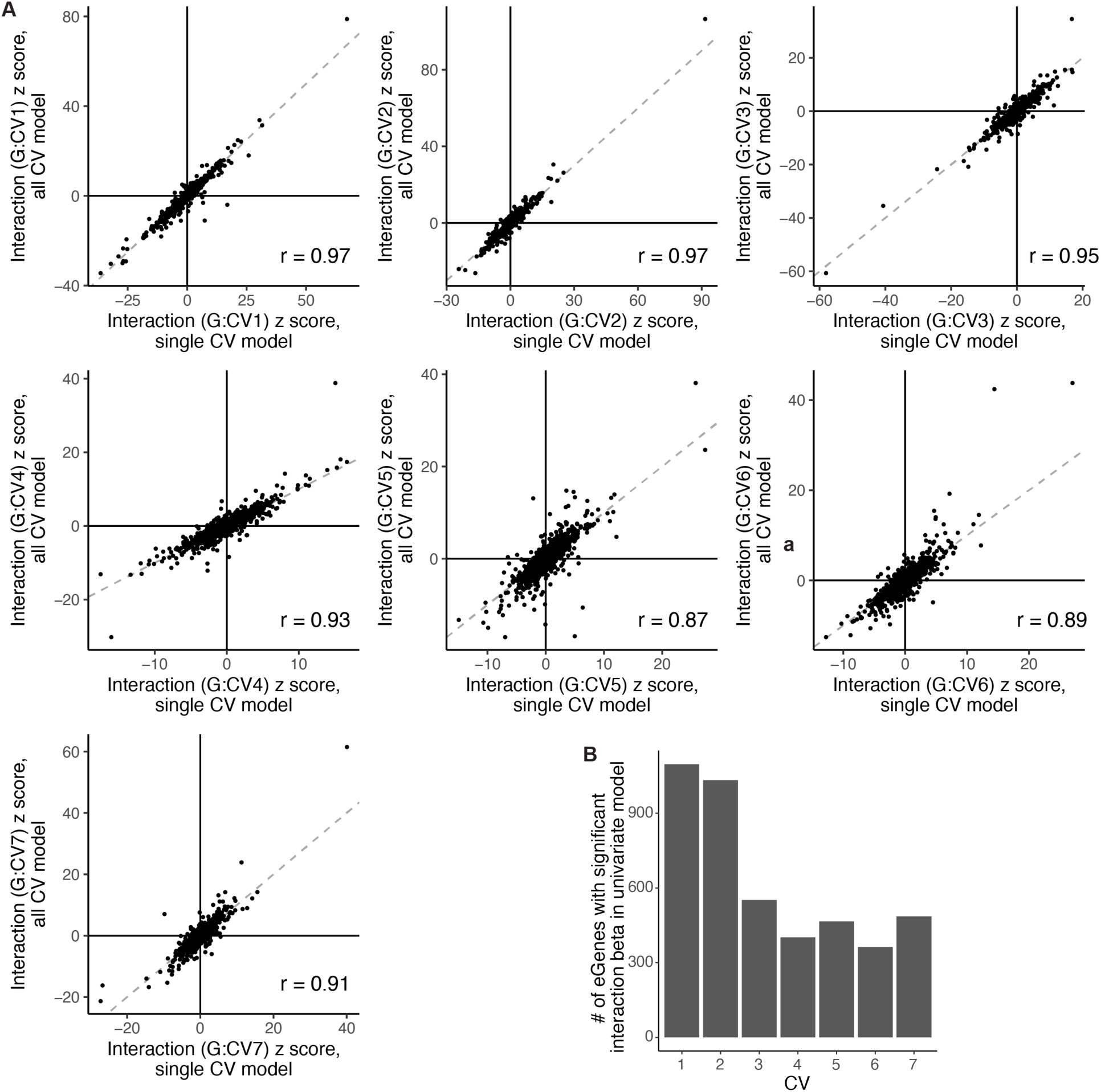
Concordance of cell-state-dependent eQTL interactions from multivariate and univariate models. **(A)** Dot plots of interaction z scores for each CV from multivariate model with 7 CVs and univariate model with one CV. Dashed line represents the identity line. r is calculated as the Pearson correlation coefficient. **(B)** Number of eGenes with significant interaction with each CV in a corresponding univariate PME model.

**Fig. S10.**
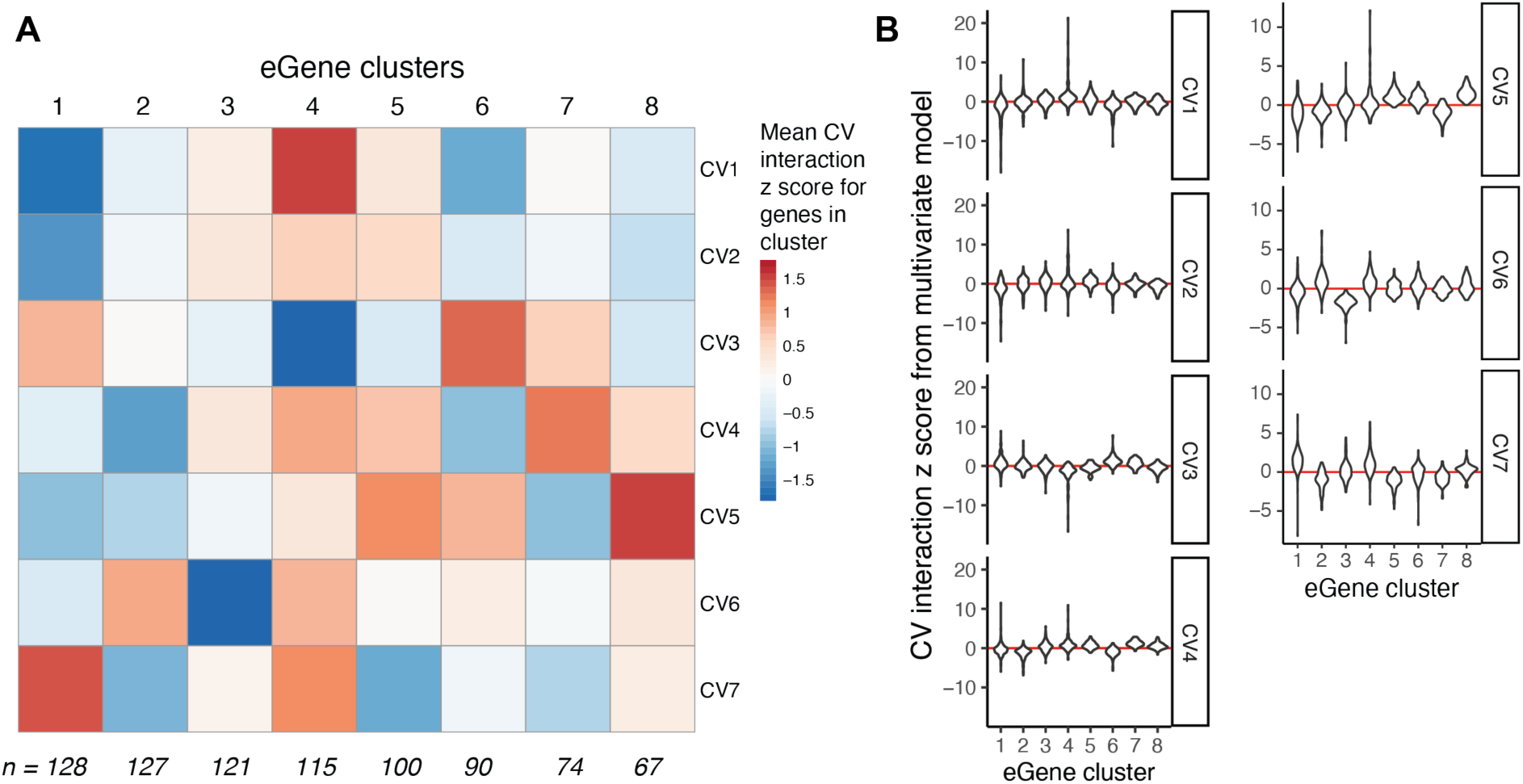
Clustering eGenes based on CV interactions. **(A)** Heatmap colored by interaction z score for each CV in the multivariate PME model, averaged acrosss eGenes in each cluster. Clusters were defined through Louvain clustering on seven interaction z scores calculated for 822 eGenes (model LRT p < .05/6511). Interaction z scores were normalized to the main genotype effect, so colors are scaled from dampening the effect (blue) to amplifying the main effect (red). **(B)** Violin plots of CV interaction z scores for eGenes in each cluster.

**Fig. S11.**
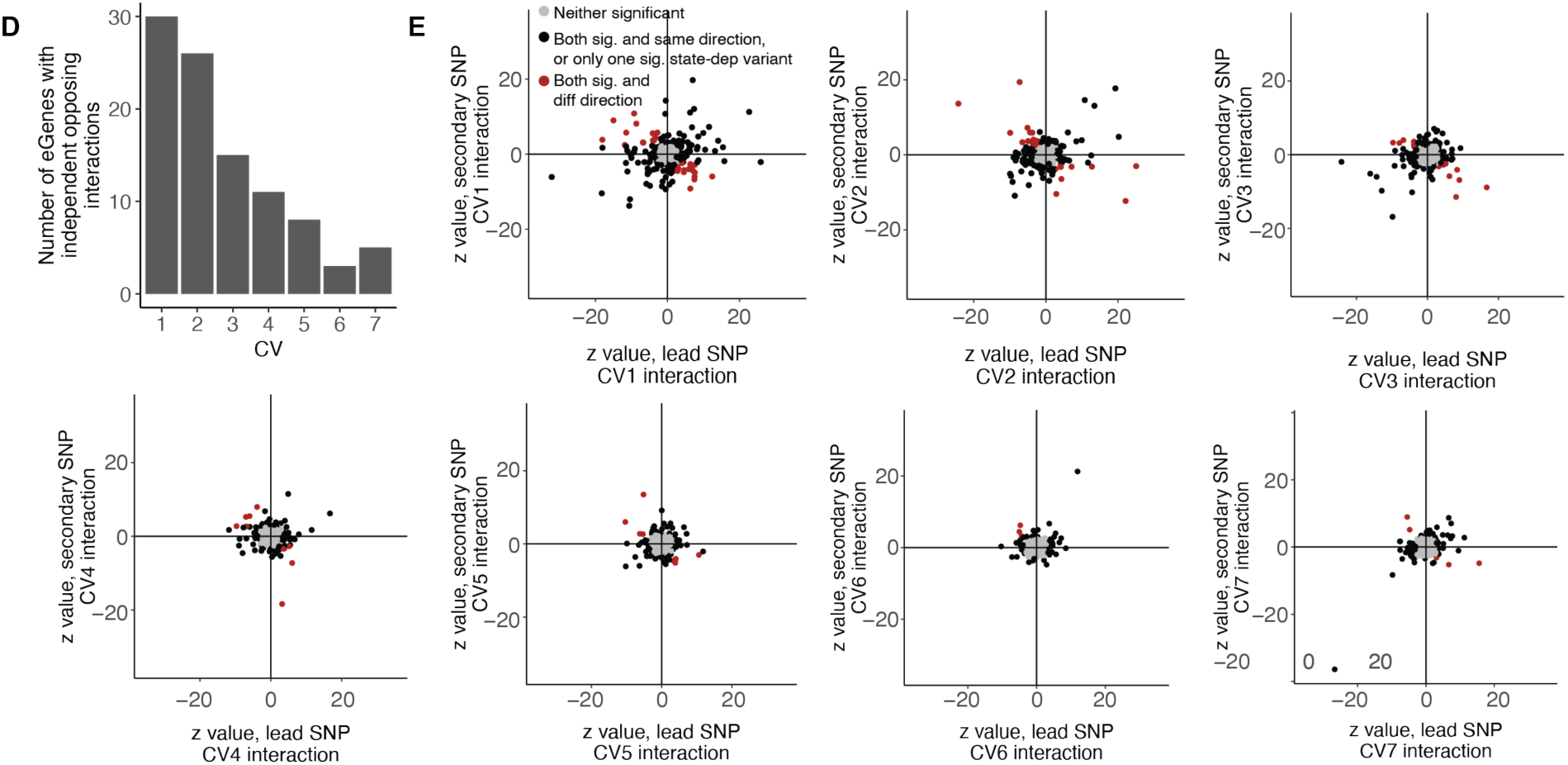
Opposite interaction directions for independent variants in a locus. **(A)** Bar plot of the number of eGenes for which the lead and secondary variants have opposite directions of interaction effect for each of the seven CVs. **(B)** Comparison of interaction effect direction for lead and secondary variants for each of 436 eGenes with 2+ independent eQTLs. Each plot corresponds to one CV, from CV1 to CV7. Each point represents an eGene. For eGenes in gray, neither lead nor secondary variant was significantly dependent on the given CV state. For eGenes in black, either only one of the two eQTLs was significantly dependent on the CV, or both were significantly state-dependent with the same direction of effect. For eGenes in red, both lead and secondary variatnts were significantly state-dependent but with different directions of interaction with the given CV.

**Fig. S12.**
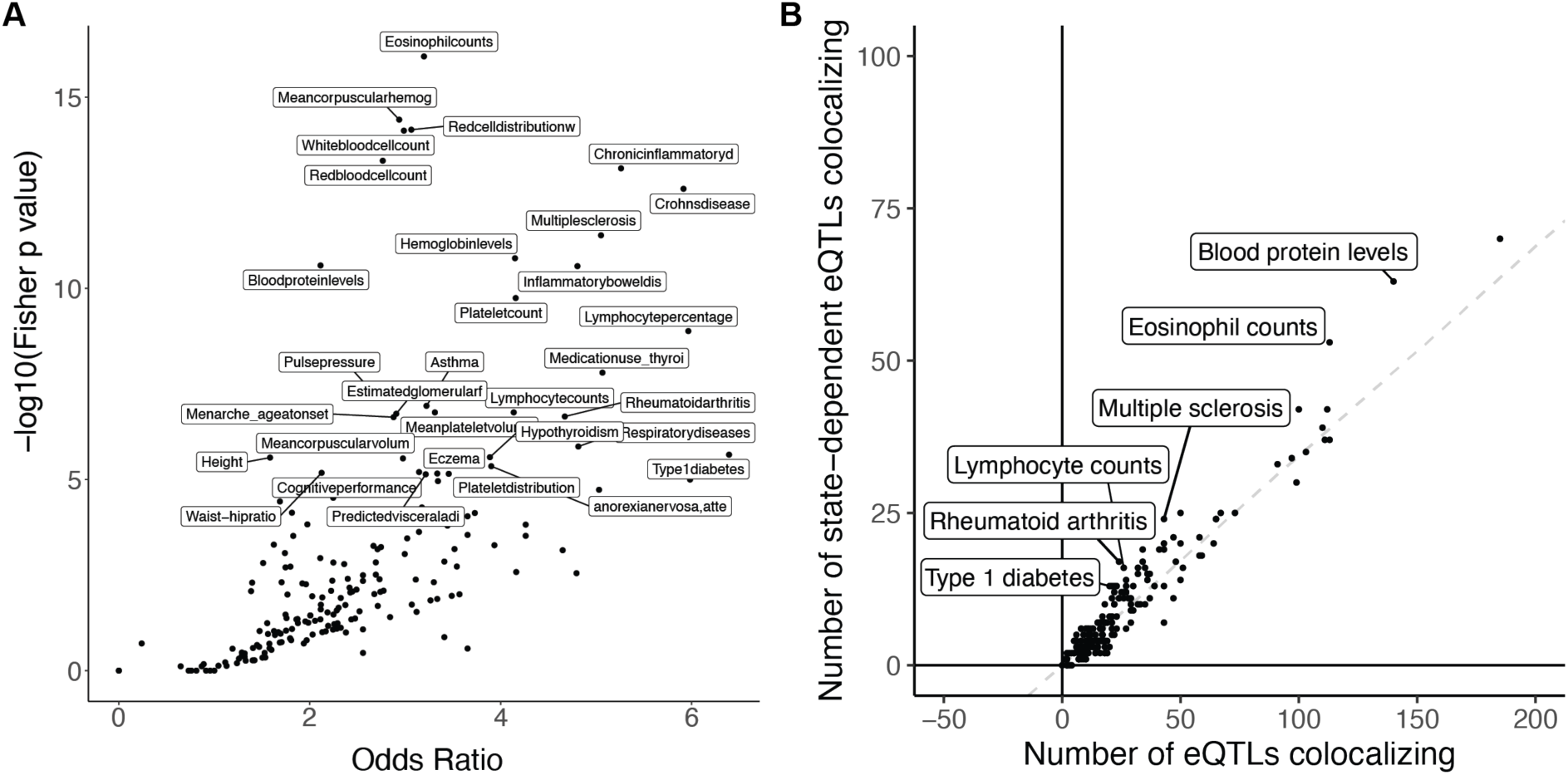
Enrichment of eQTLs in disease-associated variants. **(A)** Dot plot of traits from the GWAS catalog plotted based on the -log(Fisher p value) and odds ratio of the enrichment test comparing the proportion of GWAS variants colocalizing with memory-T cell eQTLs for one trait compared to all other traits. Labeled traits have p < 10^-5^. **(B)** Dot plot of traits from the GWAS catalog plotted based on the number of GWAS variants colocalizing with state-dependent eQTLs compared to the total number of GWAS variants colocalizing with eQTLs. The dashed line represents the overall proportion of state-dependent eQTLs (2,237/6,511 = 0.34) and labeled traits have Fisher p < 0.01.

**Fig. S13.**
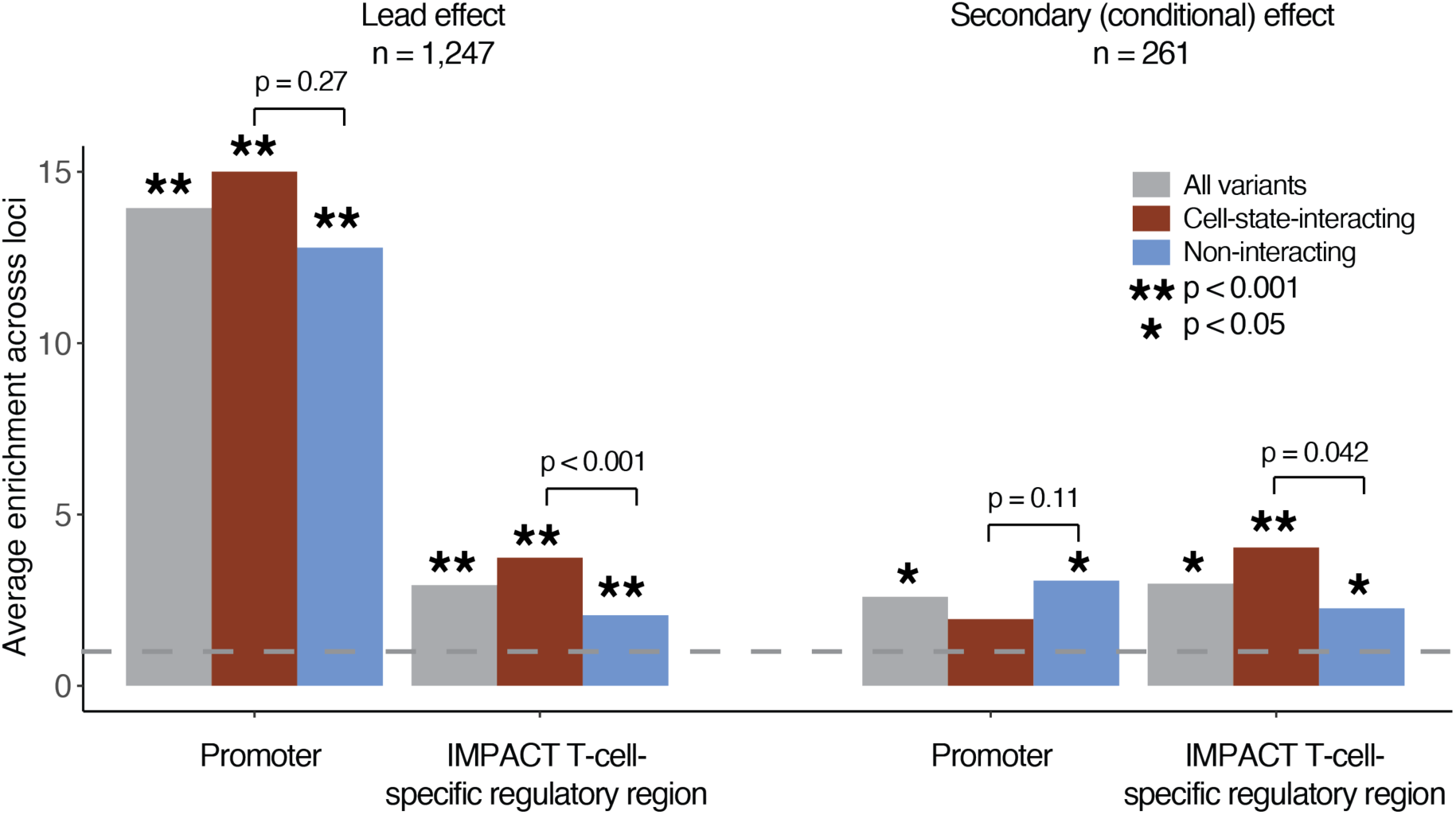
Regulatory region enrichment of eQTL effects from multi-ancestry analysis of European (BLUEPRINT) and Peruvian data. We calculated the enrichment of lead effects or independent secondary (conditional) effects in promoter or T-cell-specific regulatory regions. Analysis was limited to loci that were also significant eGenes in Peruvian analysis and where at least one variant had PIP >= 0.5. The height of the gray bar corresponds to the average enrichment calculated across all loci containing a variant with PIP < 0.05, red bar corresponds to the subset with significant cell-state interaction (LRT q < 0.05 in model with 7 CVs), and the blue bar corresponds to the subset without significant cell-state interaction. Bars marked with an asterisk are significant based on a one-sided permutation p value. Each pair of interacting/non-interacting bars is labeled with a one-sided permutation p value for the difference (interacting minus non-interacting). The gray dotted line indicates enrichment statistic = 1.

## Captions for Tables S1-23

**Table S1. Pseudobulk memory T cell eQTLs in a Peruvian cohort.** (See Nathan_etal_SuppTables.xlsx, tab 1) Lead eQTL variant for each eGene tested in a linear model of pseudobulk gene expression assayed in memory T cells from a Peruvian cohort. All SNP coordinates are from GRCh38. The model adjusts for donor’s age and sex, 5 genotype PCs, and 45 PEER factors. P values are from FastQTL beta approximation-based permutation.

**Table S2. Ancestry-specific eQTL variants.** (See Nathan_etal_SuppTables.xlsx, tab 2) eQTL variants in the Peruvian dataset driven by variants that are rare (MAF < 0.05) in 1KG EUR population.

**Table S3. Conditional pseudobulk memory T cell eQTLs.** (See Nathan_etal_SuppTables.xlsx, tab 3) Secondary eQTL variant for each eGene tested in a linear model of pseudobulk gene expression assayed in memory T cells from a Peruvian cohort, after regressing out the lead effect. The model adjusts for donor’s age and sex, 5 genotype PCs, and 45 PEER factors. P values are from FastQTL beta approximation-based permutation.

**Table S4. Gene set enrichment for loadings on CVs 1-3** (See Nathan_etal_SuppTables.xlsx, tab 4) Top 10 gene sets enriched for CVs1-3 based on genes’ loadings on each CV. P values and enrichment statistics are from the fgsea R package.

**Table S5. Average CV scores by memory T cell cluster** (See Nathan_etal_SuppTables.xlsx, tab 5) Average score along each CV for cells in each cluster. Clusters were defined in Nathan, et al. by projecting cells into a low-dimensional embedding based on CCA of paired mRNA and surface protein, constructing a shared nearest neighbor graph, and conducting Louvain clustering at resolution = 2. Clusters were annotated based on differentially expressed genes and proteins.

**Table S6. Single-cell Poisson model of memory T cell eQTLs.** (See Nathan_etal_SuppTables.xlsx, tab 6) eQTL effects calculated with the PME model (without cell state interactions) for significant eGenes and lead variants identified in the pseudobulk analysis. The model adjusts for donor’s age and sex, percent MT UMIs and number of UMIs per cell, 5 genotype PCs, 5 expression PCs, and has random effects for donor and library preparation pool. P values are from an LRT comparing the model with and without the genotype term.

**Table S7. Single-cell Poisson model of memory T cell eQTLs’ interaction with prior TB status.** (See Nathan_etal_SuppTables.xlsx, tab 7) eQTL interactions with donors’ prior TB progression status calculated with the PME model for significant eGenes and lead variants identified in the pseudobulk analysis. The model adjusts for donor’s age and sex, percent MT UMIs and number of UMIs per cell, 5 genotype PCs, 5 expression PCs, and has random effects for donor and library preparation pool. P values are from an LRT comparing the model with and without the TB status term.

**Table S8. Single-cell Poisson model of memory T cell eQTLs’ dependence on CD4+ state.** (See Nathan_etal_SuppTables.xlsx, tab 8) eQTL interactions with cells’ CD4+ state calculated with the PME model for significant eGenes and lead variants identified in the pseudobulk analysis. CD4+ cells were defined based on normalized surface protein expression measured in CITE-seq (CD4+CD8-). The model adjusts for donor’s age and sex, percent MT UMIs and number of UMIs per cell, 5 genotype PCs, 5 expression PCs, and has random effects for donor and library preparation pool. P values are from an LRT comparing the model with and without the CD4+ state interaction term.

**Table S9. Pseudobulk memory T cell eQTLs in CD4+ cells only.** (See Nathan_etal_SuppTables.xlsx, tab 9) eQTLs in CD4+ memory T cells calculated with a pseudobulk linear model in FastQTL for significant eGenes and lead variants identified in the pseudobulk analysis. CD4+ cells were defined based on normalized surface protein expression measured in CITE-seq (CD4+CD8-). The model adjusts for donor’s age and sex, 5 genotype PCs, and 45 PEER factors and we calculated a nominal P value for the genotype effect.

**Table S10. Single-cell Poisson model of memory T cell eQTLs in CD4+ cells only.** (See Nathan_etal_SuppTables.xlsx, tab 10) eQTL in CD4+ memory T cells calculated with the PME model for significant eGenes and lead variants identified in the pseudobulk analysis. CD4+ cells were defined based on normalized surface protein expression measured in CITE-seq (CD4+CD8-). The model adjusts for donor’s age and sex, percent MT UMIs and number of UMIs per cell, 5 genotype PCs, 5 expression PCs, and has random effects for donor and library preparation pool. P values are from an LRT comparing the model with and without genotype term.

**Table S11. Single-cell linear model of memory T cell eQTLs.** (See Nathan_etal_SuppTables.xlsx, tab 11) eQTL effects calculated with the LME model (without cell state interactions) for significant eGenes and lead variants identified in the pseudobulk analysis. The model adjusts for donor’s age and sex, percent MT UMIs and number of UMIs per cell, 5 genotype PCs, 5 expression PCs, and has random effects for donor and library preparation pool. P values are from an LRT comparing the model with and without the genotype term.

**Table S12. Single-cell Poisson model of memory T cell eQTLs’ dependence on CV1.** (See Nathan_etal_SuppTables.xlsx, tab 12) eQTL interactions with cells’ CV1 calculated with the PME model for significant eGenes and lead variants identified in the pseudobulk analysis. The model adjusts for donor’s age and sex, percent MT UMIs and number of UMIs per cell, 5 genotype PCs, 5 expression PCs, and has random effects for donor and library preparation pool. P values are from an LRT comparing the model with and without the CV1 interaction term.

**Table S13. Single-cell Poisson model of memory T cell eQTLs’ dependence on Treg state.** (See Nathan_etal_SuppTables.xlsx, tab 13) eQTL interactions with Tregs calculated with the PME model for significant eGenes and lead variants identified in the pseudobulk analysis. The Tregs were identified through clustering in Nathan, et al. The model adjusts for donor’s age and sex, percent MT UMIs and number of UMIs per cell, 5 genotype PCs, 5 expression PCs, and has random effects for donor and library preparation pool. P values are from an LRT comparing the model with and without the Treg interaction term.

**Table S14. Single-cell Poisson model of memory T cell eQTLs’ dependence on CVs 1-7.** (See Nathan_etal_SuppTables.xlsx, tab 14) eQTL interactions with CVs1-7 calculated with the PME model for significant eGenes and lead variants identified in the pseudobulk analysis. The model adjusts for donor’s age and sex, percent MT UMIs and number of UMIs per cell, 5 genotype PCs, 5 expression PCs, and has random effects for donor and library preparation pool. P values are from an LRT comparing the model with and without the CV1-7 interaction terms.

**Table S15. Single-cell Poisson model of memory T cell eQTLs’ dependence on CV2.** (See Nathan_etal_SuppTables.xlsx, tab 15) eQTL interactions with cells’ CV2 calculated with the PME model for significant eGenes and lead variants identified in the pseudobulk analysis. The model adjusts for donor’s age and sex, percent MT UMIs and number of UMIs per cell, 5 genotype PCs, 5 expression PCs, and has random effects for donor and library preparation pool. P values are from an LRT comparing the model with and without the CV2 interaction term.

**Table S16. Single-cell Poisson model of memory T cell eQTLs’ dependence on CV3.** (See Nathan_etal_SuppTables.xlsx, tab 16) eQTL interactions with cells’ CV3 calculated with the PME model for significant eGenes and lead variants identified in the pseudobulk analysis. The model adjusts for donor’s age and sex, percent MT UMIs and number of UMIs per cell, 5 genotype PCs, 5 expression PCs, and has random effects for donor and library preparation pool. P values are from an LRT comparing the model with and without the CV3 interaction term.

**Table S17. Single-cell Poisson model of memory T cell eQTLs’ dependence on CV4.** (See Nathan_etal_SuppTables.xlsx, tab 17) eQTL interactions with cells’ CV4 calculated with the PME model for significant eGenes and lead variants identified in the pseudobulk analysis. The model adjusts for donor’s age and sex, percent MT UMIs and number of UMIs per cell, 5 genotype PCs, 5 expression PCs, and has random effects for donor and library preparation pool. P values are from an LRT comparing the model with and without the CV4 interaction term.

**Table S18. Single-cell Poisson model of memory T cell eQTLs’ dependence on CV5.** (See Nathan_etal_SuppTables.xlsx, tab 18) eQTL interactions with cells’ CV5 calculated with the PME model for significant eGenes and lead variants identified in the pseudobulk analysis. The model adjusts for donor’s age and sex, percent MT UMIs and number of UMIs per cell, 5 genotype PCs, 5 expression PCs, and has random effects for donor and library preparation pool. P values are from an LRT comparing the model with and without the CV5 interaction term.

**Table S19. Single-cell Poisson model of memory T cell eQTLs’ dependence on CV6.** (See Nathan_etal_SuppTables.xlsx, tab 19) eQTL interactions with cells’ CV6 calculated with the PME model for significant eGenes and lead variants identified in the pseudobulk analysis. The model adjusts for donor’s age and sex, percent MT UMIs and number of UMIs per cell, 5 genotype PCs, 5 expression PCs, and has random effects for donor and library preparation pool. P values are from an LRT comparing the model with and without the CV6 interaction term.

**Table S20. Single-cell Poisson model of memory T cell eQTLs’ dependence on CV7.** (See Nathan_etal_SuppTables.xlsx, tab 20) eQTL interactions with cells’ CV7 calculated with the PME model for significant eGenes and lead variants identified in the pseudobulk analysis. The model adjusts for donor’s age and sex, percent MT UMIs and number of UMIs per cell, 5 genotype PCs, 5 expression PCs, and has random effects for donor and library preparation pool. P values are from an LRT comparing the model with and without the CV7 interaction term.

**Table S21. GO Term enrichment in eGene clusters.** (See Nathan_etal_SuppTables.xlsx, tab 21) Top 5 Gene Ontology term gene sets enriched for overlap with eGenes in each of the eight eGene clusters. These clusters were defined through Louvain clustering on the z scores for each eGene’s interactions with each of the 7 CVs, with signs corrected to be relative to the main genotype effect for that eGene. P values are from a one-sided Fisher test.

**Table S22. eQTL enrichment among GWAS variants by trait.** (See Nathan_etal_SuppTables.xlsx, tab 22) Enrichments of memory T cell eQTLs in variants associated with traits in the GWAS Catalog. Variants were considered to be overlapping if they had r^2^ > .5 in both 1KG EUR and PEL populations. P values are from a Fisher test comparing the proportion of eQTLs in a trait’s GWAS variants to the proportion of eQTLs in GWAS variants for all other GWAS Catalog traits.

**Table S23. State-dependent eQTL enrichment among GWAS variants by trait.** (See Nathan_etal_SuppTables.xlsx, tab 23) Enrichments of state-dependent memory T cell eQTLs in variants associated with traits in the GWAS Catalog. Variants were considered to be overlapping if they had r^2^ > .5 in both 1KG EUR and PEL populations. P values are from a Fisher test comparing the proportion of state-dependent eQTLs in a trait’s GWAS variants to the proportion of non-state-dependent eQTLs in GWAS variants for that trait.

## Acknowledgment

### Funding

National Institutes of Health grant U19AI111224 (SR, DBM, MM)

National Institutes of Health grant UH2AR067677 (SR)

National Institutes of Health grant T32HG002295 (AN)

National Institutes of Health grant T32AR007530 (AN)

National Institutes of Health grant U01HG009379 (SR)

National Institutes of Health grant R01AI049313 (DBM)

National Institutes of Health grant R01AR063759 (SR)

## Author contributions

Conceptualization: AN, SR

Methodology: AN, SR, AP

Formal analysis: AN, SA, KI, TA, YL

Investigation: JIB, YB, SS Resources: LL, MBM

Supervision: SR, DBM, MBM Writing – original draft: AN, SR

Writing – review & editing: all authors

## Data and materials availability

All data are available in the main text or the supplementary materials. Code will be available on Github upon acceptance.

